# Structural variation shapes regulatory and evolutionary diversity at the HLA locus

**DOI:** 10.64898/2026.04.23.720469

**Authors:** Aleix Canalda-Baltrons, Matthew Silcocks, Philippa M. Saunders, Sarah F. Jackson, Lachlan J.M. Coin, Sarah J. Dunstan

**Affiliations:** Department of Infectious Diseases, The University of Melbourne at the Peter Doherty Institute for Infection and Immunity, Melbourne, VIC, Australia; Department of Microbiology and Immunology, The University of Melbourne at the Peter Doherty Institute for Infection and Immunity, Melbourne, VIC, Australia; National Centre for Indigenous Genomics, John Curtin School of Medical Research, Australian National University, Canberra, ACT, Australia

## Abstract

The human leukocyte antigen (HLA) region is among the most polymorphic loci in the human genome and plays a central role in immune function, yet the contribution of structural variation to its genetic and regulatory diversity remains poorly characterised. Using 460 phased, near-complete human genome assemblies from globally diverse populations, we systematically mapped structural variation and gene content across the HLA locus. We show that the HLA region contains substantially more structural variation than any other region of chromosome 6. At the HLA class II locus, all individuals could be assigned to one of 13 distinct HLA-DR-DQ structural haplotypes, whereas the *HLA-A* region comprised four major haplotypes, which we found to be interspersed among non-human primate lineages. These structural haplotypes exhibit marked differences in population frequency and show increasing allelic diversity over European prehistory. Integration of Iso-Seq and RNA-Seq data revealed that structural haplotypes are associated with differences in HLA gene expression, suggesting that structural variation directly influences immune gene regulation. Together, our results identify structural variation as a key and previously underappreciated contributor to HLA regulatory diversity, with broad functional and evolutionary implications for human immunity.

**Manuscript summary:** Structural variation drives HLA haplotype diversity and gene expression differences across global human populations.

## 1 Introduction

The human leukocyte antigen (HLA) region on chromosome 6 encodes the most polymorphic genes in the human genome and is central to adaptive immunity. Classical HLA class I and II molecules present peptide antigens to T cells, shaping immune responses to pathogens, vaccines, and cancer, while also influencing autoimmune disease risk (*1,2*). This functional breadth is supported by exceptional genetic diversity, with over 42,000 allelic variants catalogued (*3*), concentrated particularly in antigen-binding regions that modulate peptide specificity and disease susceptibility (*3, 4*).

Despite its importance, structural variation (SV) in the HLA region remains poorly characterised due to complex duplication and repetitive sequences that hinder detection. Recent advances from the Human Pangenome Reference Consortium (HPRC) provide haplotype-resolved assemblies and, coupled with expression data (Iso-Seq, RNA-Seq), enable systematic SV analysis (*5, 6*). While non-coding variants like SNPs affect HLA gene regulation, the contribution of SVs to expression variation is largely unknown, despite growing evidence that gene expression levels, especially in class II genes, influence immune phenotypes, such as vaccine responses (*7–9*).

The HLA locus has been shaped by long-term balancing selection that maintains high allelic diversity in populations, enhancing resistance to a broad range of pathogens, while also incurring increased risk of autoimmune disease (*10, 11*). Some structural haplotypes, notably around *HLA-DRB1*, predate great ape speciation (*12*), but the evolutionary dynamics of these haplotypes in recent ancient time remain unclear.

Here, we analyze 460 fully phased HLA assemblies from diverse populations to map structural variation and characterize its impact on gene expression. We identify SV-expression quantitative trait locis (SV-eQTLs) influencing HLA expression, reveal major expression differences between HLA-DR-DQ structural haplotypes, and clarify evolutionary origins with ape genome comparisons. Using ancient DNA, we trace historical shifts in HLA-DR-DQ haplotypes. Our findings establish structural variation as a key determinant of HLA regulatory diversity, linking genomic architecture to immune function and evolution.

## 2 Results

### 2.1 Structural variability across the HLA locus in 460 near-T2T assemblies

The HPRC Release 2 assemblies had a total of 309 T2T chromosome 6 assemblies, with the remaining 155 assemblies having between 2-15 contigs with assembly lengths ranging from 169 Mb to 178 Mb (CHM13: 172,126,628) (figure S1a). After removing four assemblies with gaps across the HLA region, we were left with a final set of 460 fully-resolved HLA regions for analysis from diverse ancestries worldwide (Fig.1a). Notably, longer HLA haplotype sequences contained higher repeat content (figure S1b). We annotated 27-33 HLA genes per haplotype, consistent with previous studies (*13*), and found that the number of non-HLA genes and pseudogenes ranged from 139 to 146. Interestingly, we observed an inverse relationship between the number of HLA genes and non-HLA genes in African populations (figure S1c-d). Querying the IPD-IMGT/HLA database (v3.55) using Immuannot (*14*) revealed 1,914 alleles across the HLA genes not previously reported, including 190 in *HLA-DRB1*, 22 in *HLA-DQA1* and 12 in *HLA-DQB1*; class II genes which have been associated with autoimmune and infectious disease susceptibility (*4, 11, 15–18*) (table S1). Furthermore, our data highlight the under-representation of non-European populations in the HLA database, as evidenced by the comparatively lower percentage of newly annotated alleles found in European haplotypes (figure S2).

**Figure 1.**
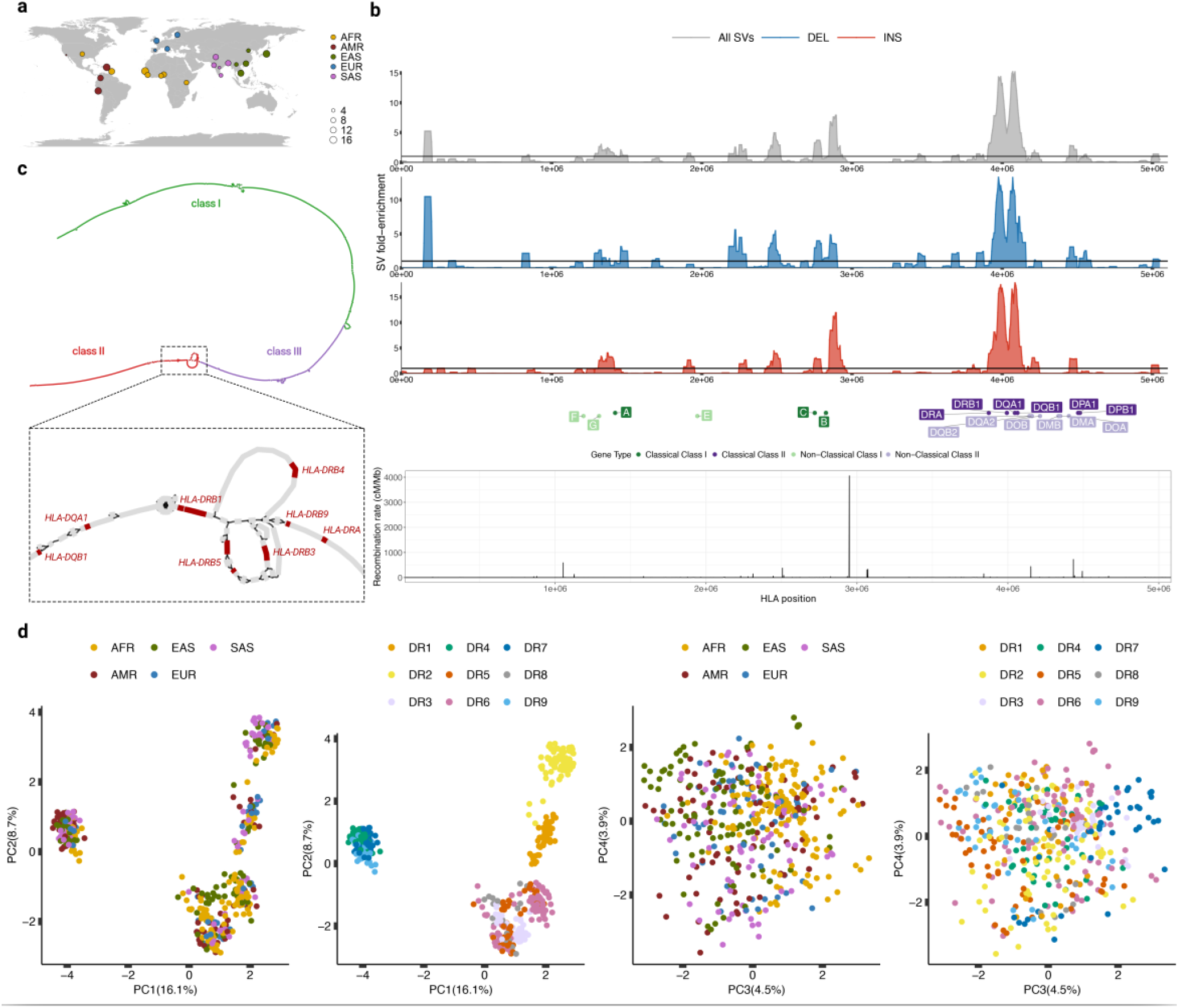
Detecting structural variation across the HLA locus. **a** Map showing the geographical location of the populations included in our analysis and their sample sizes. **b** Distribution of SVs across the HLA locus by their fold-enrichment relative to the rest of the HLA region across 50 kb windows and 5 kb steps, segregated by SV type across all haplotypes. Horizontal lines represent no fold change of a specific position from the rest of the region. The classical and non-classical functional HLA genes are annotated. The bottom plot shows the recombination rates detected across the HLA region. **c** HLA minigraph graph representation, colored by the three HLA classes, zooming into the *HLA-DRB1* structurally variable locus. **d** PCA plots of HLA SV genotypes for each haplotype. DR group determined by typed *HLA-DRB1* alleles.

We identified a total of 519 unique structural variants (SVs) across the HLA locus, with an average of 68 SVs per haplotype when mapped to the CHM13 reference genome. Compared to the Human Genome Structural Variation Consortium phase 3 (HGSVC3) (*13*) HLA SV dataset, we found 136 novel SVs, while 85 were HGSVC3-unique, most being singletons. A prominent SV hotspot was found across the HLA-DR-DQ region, consistent with the high repetitive content found within it (*13*) and the relatively low recombination rates (Fig.1b). A pangenome reference graph representation of the structural variation found across the HLA locus confirmed the prominent hotspots for SVs across the HLA genes and pseudogenes in both classes I and II, as well as the region surrounding *C2, C4A* and *C4B* in class III. The graph was also able to capture different classical HLA-DR haplotypes (Fig.1c). HLA-DR haplotypes have been commonly classified into 9 groups comprising 5 main structural haplotypes, traditionally defined by their *HLA-DRB1* allele and associated gene and pseudogene content, using the DR group system. Previous population-scale studies have shown that these DR groups capture the major structural configurations of the HLA-DR locus observed in humans (*13*). A principal component analysis (PCA) confirmed that structural variation across the whole HLA locus was predominantly driven by classical HLA-DR groups, typed using *HLA-DRB1* alleles, and not by continental populations (Fig.1d). We found 10 individuals (0.2%) with a full *MICA* deletion, from Africa, East Asia and America, a frequency concordant with previous results (*19*). Other full gene deletions pertained to *HLA-W, HLA-DPA2* at low frequencies (in 2 and 1 individuals, respectively), as well as *HLA-H, -T, -K* and *-U* with higher frequencies (13%). However, the gene with the highest frequency of SVs across haplotypes was *HLA-DQA1* (figure S3). Structural variant calls were also validated using Hifi long read datasets (Supplementary Text 1; Supplementary Methods 1).

These findings underscore the high structural variability within HLA class II compared to the other classes, emphasizing the need for a more comprehensive characterisation of complete haplotypes across individuals from diverse backgrounds. Given the strong association of HLA class I genes with infectious disease susceptibility, cancer and organ transplant (*17*), we also sought to characterise the structural haplotypes of HLA class I genes in our dataset.

### 2.2 HLA structural genomic diversity across human populations

The global frequencies of the identified SVs and described haplotypes may contribute to the differential susceptibility to infectious and autoimmune diseases. When grouping HLA-DR haplotypes using the classical DR group system (DR1, DR2, DR3, etc.), we found that DR3/5/6 was the overall most frequent haplotype, especially in African populations (55%), while DR4/7/9 was the most frequent haplotype in American populations (figure S4).

We also aimed to understand whether HLA SV alleles were mostly shared across populations or unique to them. We found that an average of 65% of structural variation across the HLA locus was shared among all continental populations (range 53.8%-81.3%; figure S5a). Previous studies found lower global SV sharing genome-wide (*20*), suggesting most structural variation across the HLA locus is ancient and has been maintained across populations due to balancing selection. To further characterise the structural variation among diverse populations, we leveraged long-read sequencing datasets from 976 individuals from the 1,000 Genomes Project (*20*). We found a total of 11,908 SVs across chr6 of which 441 belonged to the HLA region. This showed that the 460 phased assemblies yielded more SVs than 976 ONT-sequenced individuals across the same region. We found that the HLA region is significantly more SV rich than the rest of chr6 (two-sided t-test; *p*-value < 2e-16), across all populations (figure S5b). Interestingly, despite African populations having a significantly higher density of SVs across chr6 (figure S5b, figure S6a; two-sided t-test; *p*-value < 2e-16), it was not significantly higher across HLA (figure S5b, figure S6b; two-sided t-test; *p*-value = 0.88). This could have been driven by various factors such as balancing selection or archaic introgression, which served to further diversify the HLA region in other populations (*21*). Finally, we wanted to determine whether the HLA was enriched for rare SVs (higher proportion of singletons) compared to a random, 5 Mb region also in chr6. We found that 16% of the SVs across the HLA were singletons, while 26.7% for the non-HLA region, indicating that the HLA was not enriched in rare structural variation (figure S5c).

### 2.3 Structural haplotypes across the HLA-DR-DQ region

Our HLA assemblies include representatives of all known DR groups, with sample counts ranging from 25 (DR8) to 93 (DR6). However, these classifications have largely considered HLA-DR in isolation. Because HLA-DR and HLA-DQ form a tightly linked genomic region (Supplementary Text 2; table S2), jointly analysing their structural variation provides an opportunity to refine these haplotypes and better understand their evolutionary relationships.

To refine existing DR group classifications across the linked HLA-DR-DQ region, we resolved the underlying structural haplotypes using a graph-based representation of the locus. By mapping each assembled haplotype onto the HLA graph (see Methods), we identified the distinct genomic paths that define each structure. Although individual DR groups initially appeared to contain many haplotypes—ranging from 3 in DR1 to 28 in DR6—many differed by only a few bases. Clustering DR-DQ haplotypes based on sequence similarity therefore collapsed much of this apparent diversity (Fig.2a). Within each DR group, we assigned names to haplotypes based on their relative size, designating the smallest as ‘.1’, followed by ‘.2’, ‘.3’, and so on for larger variants; this nomenclature is used throughout the manuscript. Using both within- and between-group comparisons, we ultimately resolved a total of 13 distinct structural haplotypes across HLA-DR-DQ (Fig.2b; figure S7).

**Figure 2.**
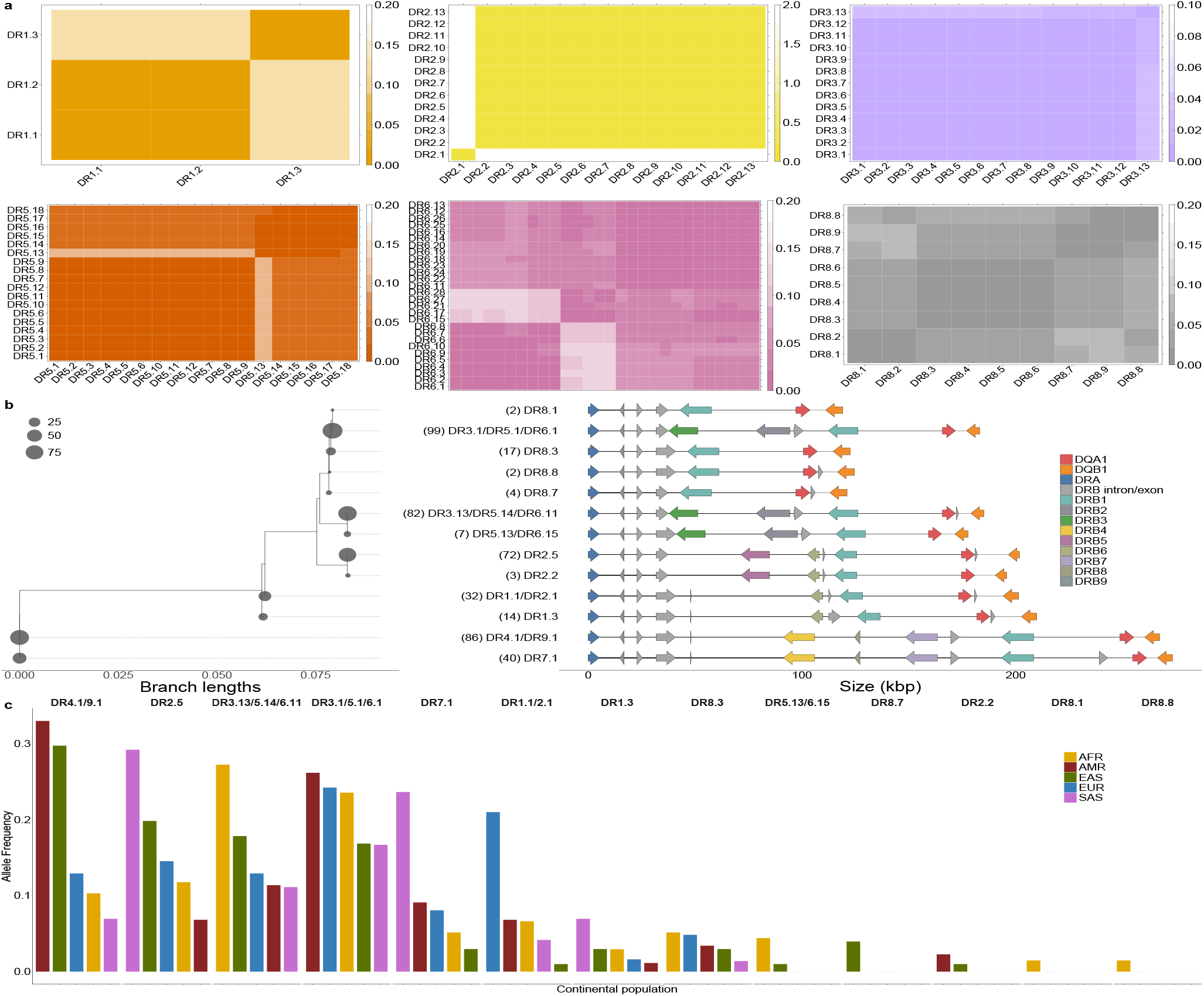
Clustering and structural diversity across HLA-DR-DQ. **a** Clustering of within-DR groups according to the distance matrix, calculated from a multiple sequence alignment. The color bar corresponds to number of different bases per 100 bp; similar haploypes will have a stronger color than dissimilar ones. **b** Thirteen distinct HLA-DR-DQ structural haplotypes identified in 460 haplotypes. The colored arrows indicate genes, while the gray arrows indicate pseudogenes. The forward strand-facing gray arrows represent solitary DRB exons/introns. The numbers in parentheses and the circle sizes indicate the number of haplotypes identified with a specific structure. Haplotypes are ordered by their relationship in the tree (left), which is generated from a multiple sequence alignment of all structures and IQtree (*58*). Consensus structures, which refer to clusters of similar structures, are indicated (right). The names of the consensus structures come from the clustering of within-group haplotypes in **a. c** HLA-DR-DQ haplotype frequency across phased assemblies from diverse populations.

Consistent with previous observations, most structural differences among these DR-DQ haplotypes were driven by the presence or absence of solitary *HLA-DRB* exons and introns, which are thought to be remnants of ancestral nonhomologous recombination events involving LINE1 and Alu elements in *HLA-DRB* intron 1 sequences (*13, 22, 23*). These features clearly distinguished several DR groups, especially DR8. A notable exception was a DR2.1 haplotype from an African individual that carried DR2 alleles at *DRB1* but shared the overall structural configuration of DR1.1, consistent with a rare recombination event between DR2 and DR1 or DR4/7/9 haplotypes (figure S8). In contrast to previous studies that focused primarily on DR, our inclusion of *DQA1* and *DQB1* allowed further refinement of these DR–DQ structural haplotypes, revealing additional solitary *DRB* exons and introns flanking these genes. For example, an *DRB* exon/intron upstream *DQA1* was present in all DR7.1 haplotypes but absent from DR4.1 and DR9.1. Finally, comparison of predicted transcription factor binding sites across the resolved structural haplotypes showed differences in both the composition (figure S9a) and number of motifs (figure S9b), suggesting that structural variation across the HLA-DR-DQ region may contribute to differential HLA gene expression.

We also sought to determine the structural variation that was driving within-DR-DQ structural haplotype diversity and where it fell. No structural diversity was detected within DR8.1, DR8.7 or DR1.3. Interestingly, we found more DRB exon/intron variations within DR-DQ structural haplotypes, not captured during the initial clustering. In DR3.1/5.1/6.1 and DR3.13/5.14/6.11 we found an extended DRB intron/exon downstream *DRB3* in 25% and 40% of cases, respectively (figure S10). Moreover, we found the DRB intron/exon upstream *DQA1*, in all DR7.1 haplotypes, was also present in 43% of DR5.13/6.15 haplotypes. We also found large variation in gene introns with two bubbles in introns 1 and 3 in *DQA1* and intron 2 in *DQB1* fixed in various haplotypes with SVs reaching up to 323 bp. Most notable, however, was the intronic variation found in *DRB1*, with an SV reaching 2,013 bp in intron 1, a likely DNA transposon excision, unique to DR2 and DR1.

Moreover, analysis of the 13 resolved DR-DQ structural haplotypes revealed substantially greater population stratification than was apparent when considering the five broad DR groups alone (figure S4). At this higher resolution, continental differentiation became markedly clearer. For example, DR4.1/9.1, whose *DRB1* alleles have been associated with protection against enteric fever (*11, 24*), is predominantly observed in American and East Asian populations (Fig.2c). In contrast, DR7.1 is highly enriched in South Asia. Similarly, DR1.1/2.1 is largely restricted to Europeans, whereas DR1.3 occurs primarily in South Asian populations. Although the broader DR8 group is most frequent in Africa, our structural classification further resolves a distinct DR8 haplotype, specific to East Asian populations. Collectively, these findings demonstrate that fine-scale structural haplotype resolution captures substantially more population-specific differentiation than classification based on the five conventional DR groups.

### 2.4 Ape evolutionary insights into the *HLA-A* region

Across the *HLA-V*-*HLA-J* region in HLA class I, we found 4 distinct structural haplotypes as described in ref. (*5*) (figure S11a). *HLA-A* alleles were tied to specific haplotypes. Specifically, the A.1 haplotype was only found in genomes with A*23/*24 alleles, A.2 was only found in two individuals with A*66 alleles, meanwhile A.3 and A.4 haplotypes shared most of the *HLA-A* alleles (figure S11a). A.3 was the most frequent across all populations, especially in European populations. The *HLA-Y*-containing A.4 was most frequent in Africans while the *HTKU-del* was most frequent in East Asian populations (figure S12) and least frequent in Africans. A pairwise alignment between A.1 and A.3 revealed the retrotransposon LINE1 as the likely origin of the *HLA-HTKU* deletion through non-allelic homologous recombination (figure S11b), which could also explain the *HLA-Y* deletion (figure S11c).

While the structural variation underlying the 5 main HLA-DR haplotypes has been extensively studied and is estimated to have started to diverge >40 million years ago (Mya) (*12, 25*), the 4 main structural haplotypes described across the *HLA-A* region, comprising HLA pseudogenes, have received less attention (*5*). The recent release of T2T ape genomes has enabled a more detailed study of the Major Histocompatibility Complex (MHC; known as HLA in humans) region in our closest relatives (*26*). Analysis of the structural haplotypes across *HLA-A* in the ape genomes could provide valuable insights into the evolutionary history of the 4 main haplotypes found in humans.

Although functional MHC genes have been annotated across the ape T2T genomes, the MHC pseudogenes across *MHC-A*, which are associated with the 4 main human structural haplotypes, remain uncharacterised. Given their association with large-scale structural variation, differences in pseudogene content could plausibly influence local genomic architecture and downstream regulatory potential. Notably, while the functional gene content is almost identical between humans and apes (*26*), the pseudogene content exhibits significant differences. A graph representation of the *MHC-A* region elucidated the structural haplotypes across the ape genomes (Fig.3a). We observed within-species structural heterogeneity in chimpanzees (PTR), gorillas (GGO) and Sumatran orangutans (PPY) whereas the other great ape species analyzed each exhibited a single structural configuration (Fig.3b). Specifically, we detected *Patr-AL* in PTR.h2, a non-classical MHC class I gene uniquely found in chimpanzees, accompanied by an *MHC-K* duplication upstream and extensive reorganization of pseudogene content. Our analysis revealed the *-H, -T, -K* and *-U* gene/pseudogene content to be highly variable. In contrast, *MHC-W* and *MHC-J* were found in nearly all haplotypes except A.2 and GGO.h1, respectively. Notably, we identified the human *HLA-Y* pseudogene as likely derived from a functional *MHC-A* orthologue in apes, featuring an 800 bp deletion that likely rendered it nonfunctional. Furthermore, we discovered an *MHC-A* pseudogene in GGO.h1, which arose from an acquired STOP codon in exon 4. These findings highlight the significant complexity and structural variability across the *MHC-A* region in apes. The *MHC-A* pseudogene annotations are provided in table S3.

**Figure 3.**
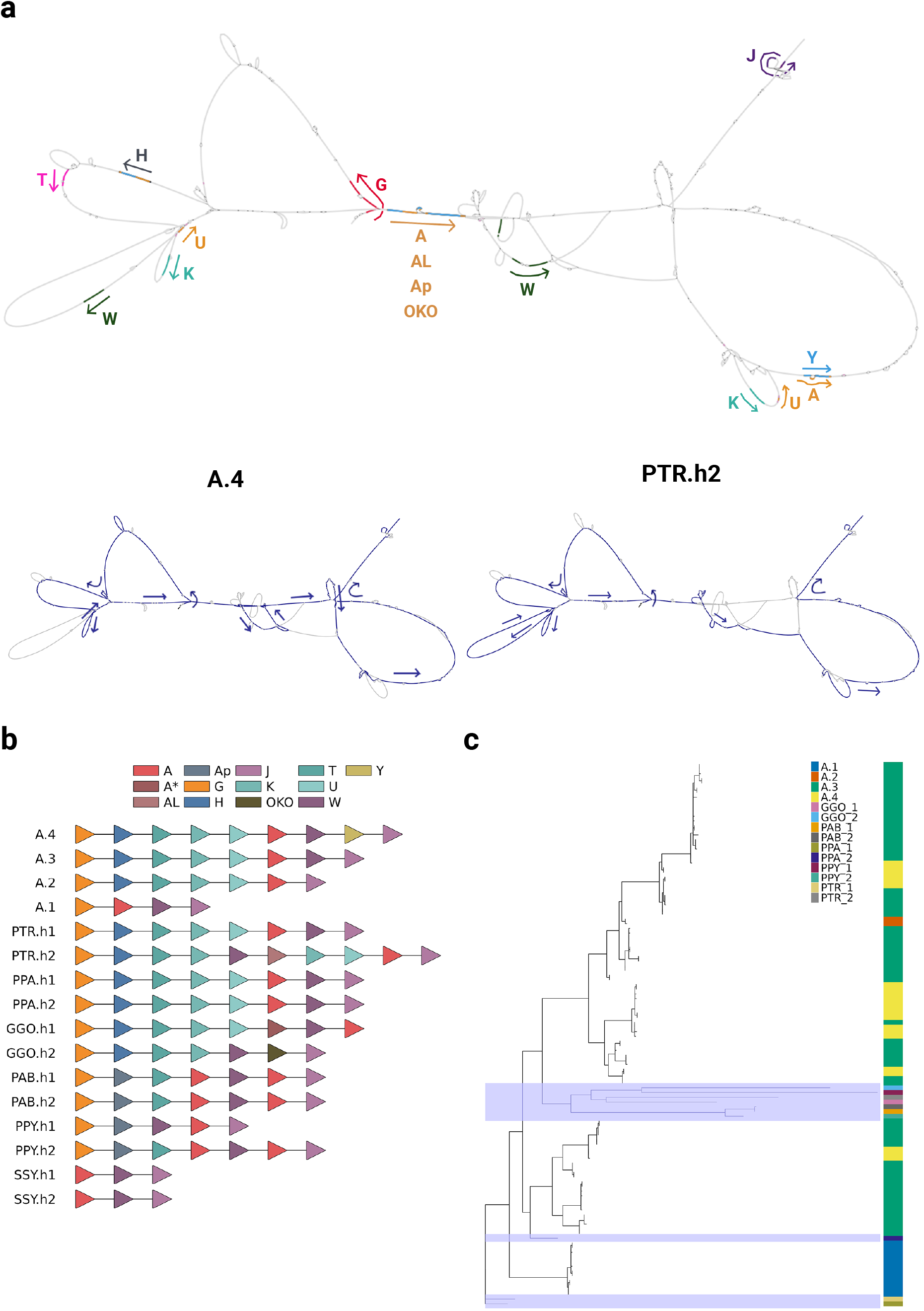
Ape structural variation across the *MHC-A*. **a** Addition of ape HLA T2T sequences to the HLA graph. Specifically, the *HLA-A* structural haplotype bubbles are shown with the direction of the genes in the graph indicated by arrows. We further show two distinct paths traversed across the HLA graph in chimpanzee haplotype 2 and human haplotype A.4, showing highly distinct structural organisation across species. **b** Gene/pseudogene content is shown across all haplotypes. AL: *Patr-AL*; A*: *MHC-A* pseudogene in GGO.h1. **c** Unrooted phylogeny of the *HLA-A* gene sequences across human and ape T2T genomes. Tiles are colored as specific structural haplotypes for each tip and the shaded blue areas highlight the ape genomes intermixed with the human genomes. Human tips that had less than 1e-6 divergence with other human branches were pruned. PTR: *Pan troglodytes*; PPA: *Pan paniscus*; GGO: *Gorilla gorilla*; PAB: *Pongo abelii*; PPY: *Pongo pygmaeus*; SSY: *Symphalangus syndactalus*.

The *HLA-A* gene flanks the 4 main human structural haplotypes (figure S11c; Fig.3b); clustering the *HLA-A* sequences across all haplotypes could reveal their evolutionary trajectories. The resulting phylogeny revealed that the A.1 haplotypes harbouring the *HTKU*-del clustered together and separately from the other human haplotypes, closer to the chimpanzee and bonobo sequences (Fig.3c). Moreover, the A.1 haplotype presents unique *HLA-A* alleles (A*23/*24), not found in other human structural haplotypes, suggestive of a distinct evolutionary trajectory. In contrast, the *HLA-Y* insertion haplotypes did not cluster together, indicating that the *HLA-Y* deletion occurred recurrently across the human phylogeny.

These results emphasise the deep evolutionary history of the *HLA-A* region, with the human lineages we characterise interspersed between Great Apes.

### 2.5 Structural variation is associated with modified HLA gene expression

To determine whether the HLA structural haplotypes as well as specific SVs have an effect on HLA gene expression, we leveraged the Iso-Seq data from 199 individuals included in the HPRC Release 2. The availability of phased assemblies for each individual enabled phasing of Iso-Seq reads, providing the resolution necessary to assess haplotype-specific gene expression within each sample. On average, across all samples and the whole HLA region, 47.7% of the reads could be phased (range of 0.22%-72.3%; Fig.4a). An initial survey across the HLA locus suggested frequent mono-allelic expression at the class III genes *C4A* and *C4B*, with approximately 40% of samples showing expression from a single haplotype (Fig.4b). After restricting to samples with fewer than 5% unphased reads, we identified 11 samples with mono-allelic expression of *C4A* and one sample with no detectable expression, as well as 39 samples with mono-allelic expression of *C4B* and four with no detectable expression. To determine whether this pattern reflected copy number variation at the C4 locus, we examined the relationship between deletions overlapping C4A and C4B and mono-allelic expression. Most deletions originated within *C4A* and extended to fully overlap *C4B*, providing a mechanistic explanation for the higher frequency of mono-allelic *C4B* expression (figure S13). Consistent with this, homozygous deletions were observed on haplotypes lacking *C4B* expression. Given the established association of *C4A* and *C4B* deficiencies with autoimmune disease susceptibility through their roles in the complement system, these results underscore the functional relevance of detecting structural variation at this locus (*27*).

**Figure 4.**
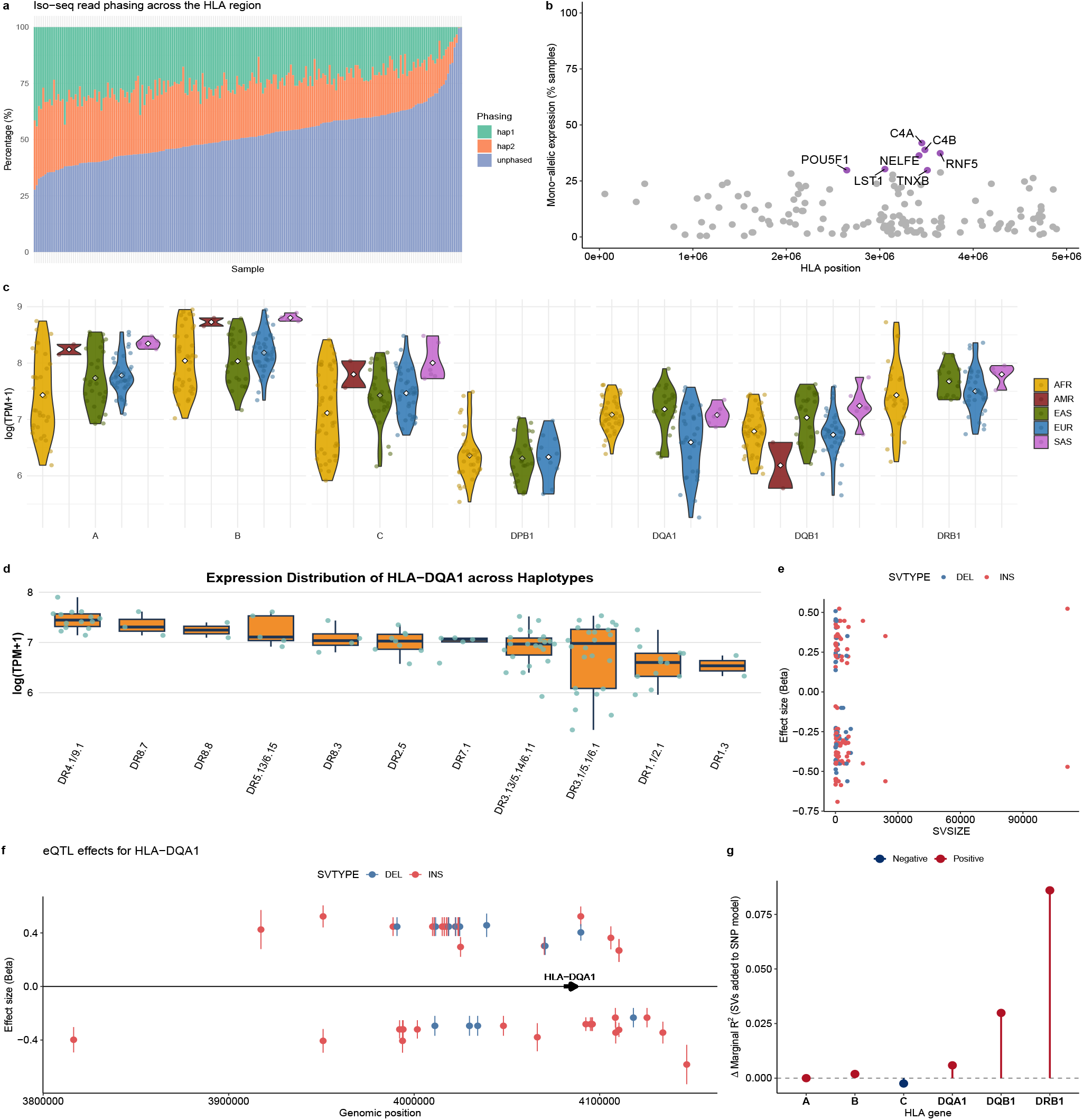
Effect of structural variation on HLA gene expression. **a** Phasing of Iso-Seq reads mapping them onto their respective phased assemblies across the whole HLA region. This is shown as a percentage from the total number of reads mapped to the HLA region. Those reads that mapped equally well to either haplotype were categorised as unphased. **b** Frequency of genes expressed in a single haplotype. Purple-colored genes represent those with mono-allelic expression in at least 30% of the samples. Here samples were not filtered out by percentage of phased reads; percentage of mono-allelic expression is overestimated. **c** HLA gene expression across continental populations as detected by RNA-Seq. A single American and 2-3 South Asian individuals were included in this analysis. The number of samples included varied across genes as only those samples that were correctly phased for that gene were included, leading to genes with no American or South Asian representation. Violin plots indicate the mean (center white dot), extending to the highest and lowest value.**d** *HLA-DQA1* expression across DR haplotypes. Box plots indicate the median (center line), interquartile range (box), and whiskers extending to 1.5 × the interquartile range. **e** Effect size of all significant SV-eQTLs, by SV size. **f** Position and effect size of significant SV-eQTLs associated with modified *HLA-DQA1* gene expression. **g** Change in marginal R^2^ of mixed-effects models when the 5 most associated structural variants (SVs) are added to models already including the 5 most associated SNPs, per HLA gene. Positive values (red) indicate increased explained variance attributable to SVs, while negative values (blue) indicate no additional explanatory power. Results highlight gene-specific contributions of structural variation to HLA expression variability.

To determine whether the Iso-Seq data could accurately quantify HLA gene expression, we compared the expression with RNA-Seq datasets from 77 samples (*6, 28–30*). We found that the expression values were not consistent across the Iso-Seq and RNA-Seq datasets for most HLA genes, especially *HLA-C*, likely due to incomplete saturation of the transcriptome in Iso-Seq data (figure S14). We therefore quantified total gene expression across the 77 samples with RNA-Seq data by aligning the reads to personalised reference genomes (see Methods). The resulting expression distributions across HLA genes were consistent with those reported in previous studies (*31*) (Fig.4c). Our approach allowed us to quantify haplotype-specific gene expression where read-phasing was successful. All HLA genes, except *HLA-DPA1* and *HLA-DPB1*, had successful phasing in at least 60% of samples (<5% unphased reads), reaching up to 81% for *HLA-C* (figure S15).

We found that HLA genes exhibited substantial variation in expression (Fig.4c). However, this variation was not uniform across populations; for example, African populations showed lower HLA class I gene expression, whereas Europeans had the lowest expression of *DQA1*. Class II differences are likely attributable to varying frequencies of structural haplotypes among populations, as the expression patterns associated with specific HLA-DR-DQ structural haplotypes were largely consistent across populations (figure S16-18). We observed that these haplotypes were associated with different expression levels of HLA class II genes (Kruskal-Wallis test, χ^2^ = 42.514, *p*-value = 1.3e-5 for *DQA1*, χ^2^ = 51.2, *p*-value = 3.8e-7 for *DQB1*, χ^2^ = 27.05, *p*-value = 2.2e-3 for *DRB1*; Fig.4d, figure S19a-b). DR3.1/5.1/6.1 presented high expression variability, likely due to greater within-haplotype structural diversity (Fig.4d, figure S10, figure S19a-b). We also sought to find unique SVs associated with HLA gene expression (expression quantitative trait loci (eQTLs)). We found 168 SV-eQTLs across the 6 classical HLA genes, though most were associated with class II (13-22 vs 3-10 independent signals for class I), indicating a stronger potential for SVs to impact class II gene expression (figure S19c-e; table S4). We found that both insertions and deletions were associated with modified gene expression, regardless of SV size (Fig.4e). A LINE1 retrotransposon insertion upstream *HLA-DQA1* was associated with a 68% increase in its expression, found in DR4.1/9.1 and DR5.13/6.15 haplotypes (Fig.4f). Four SVs downstream *HLA-DQB1* in linkage disequilibrium were the most associated with a 50% increased expression, including the DRB intron/exon sequence found between *DQA1* and *DQB1* (figure S19c). The most significant SV block associated with modified expression, however, was a linked block of 4 SVs that decreased the expression of *DRB1*, one of which was a large intron 1 deletion. Intriguingly, all SVs associated with modified *DRB1* expression result in decreased expression. This might be due to DR3.1/5.1/6.1, the haplotype found in CHM13, which exhibits one of the highest *DRB1* expressions among all haplotypes. We also found that a 1,758 bp SVA (SINE-VNTR-Alu) retrotransposon, inserted between *HLA-K* and *HLA-U*, reduced the expression of *HLA-A*, highlighting the importance of the *HLA-A* structural haplotypes. To validate these findings, we analyzed SNP eQTLs and found that they follow a similar pattern to SV-eQTLs, such as most *DQB1* eQTLs being downstream or most *DQA1* eQTLs being upstream (figure S20).

To quantify the contribution of structural variation to HLA gene expression, we compared linear mixed effects models including the top five associated SNPs for each gene with models that additionally incorporated the top five associated SVs. Inclusion of SVs resulted in gene-specific changes in explained variance. For the class II genes *HLA-DQA1, HLA-DQB1*, and *HLA-DRB1*, addition of SVs increased the variance explained (Fig.4g), indicating that these variants capture regulatory effects beyond those tagged by the strongest SNP associations. In contrast, for the class I genes *HLA-A, HLA-B*, and *HLA-C*, SVs did not highly improve model performance and in some cases slightly reduced explained variance. Together, these results indicate that structural variation contributes disproportionately to expression variability at class II HLA loci and underscore the crucial role of SVs in regulating HLA gene expression, highlighting their potential impact on immune response.

### 2.6 An increase of HLA-DR-DQ structural haplotype diversity in recent European prehistory

Given that structural variation could have phenotypic effects, including altered class II HLA gene expression, we hypothesised that the frequencies of HLA-DR-DQ structural haplotypes were shaped over time by demographic shifts and changes in pathogen load in prehistoric Europe. To test our hypothesis, we typed classical HLA genes in 1,290 ancient European genomes using OptiType (*32*). Because structural haplotypes for HLA-DR-DQ can be inferred from the first typing fields of *DRB1* and *DQA1* (Supplementary Text 3; table S5), this approach was particularly suitable. Benchmarking on simulated ancient genomes demonstrated that OptiType achieves high accuracy for first-field typing of class II HLA genes, whereas a recent pangenomic method performed poorly on similarly low-coverage, low-quality data (Supplementary Text 3; figure S21). We retained 556 genomes with a minimum coverage of 0.5× for downstream analyses. A comparison with the simulated ancient genomes revealed that those with coverage 0.5-2× had heterozygosity lower than expected, while this was not observed in genomes with >2× coverage (figure S22). In total, by keeping genomes with coverage greater than 2×, we had 168 ancient European genomes, or 336 inferred structural haplotypes, which spanned from approximately 13,600 to 600 BP (Fig.5a; table S6).

**Figure 5.**
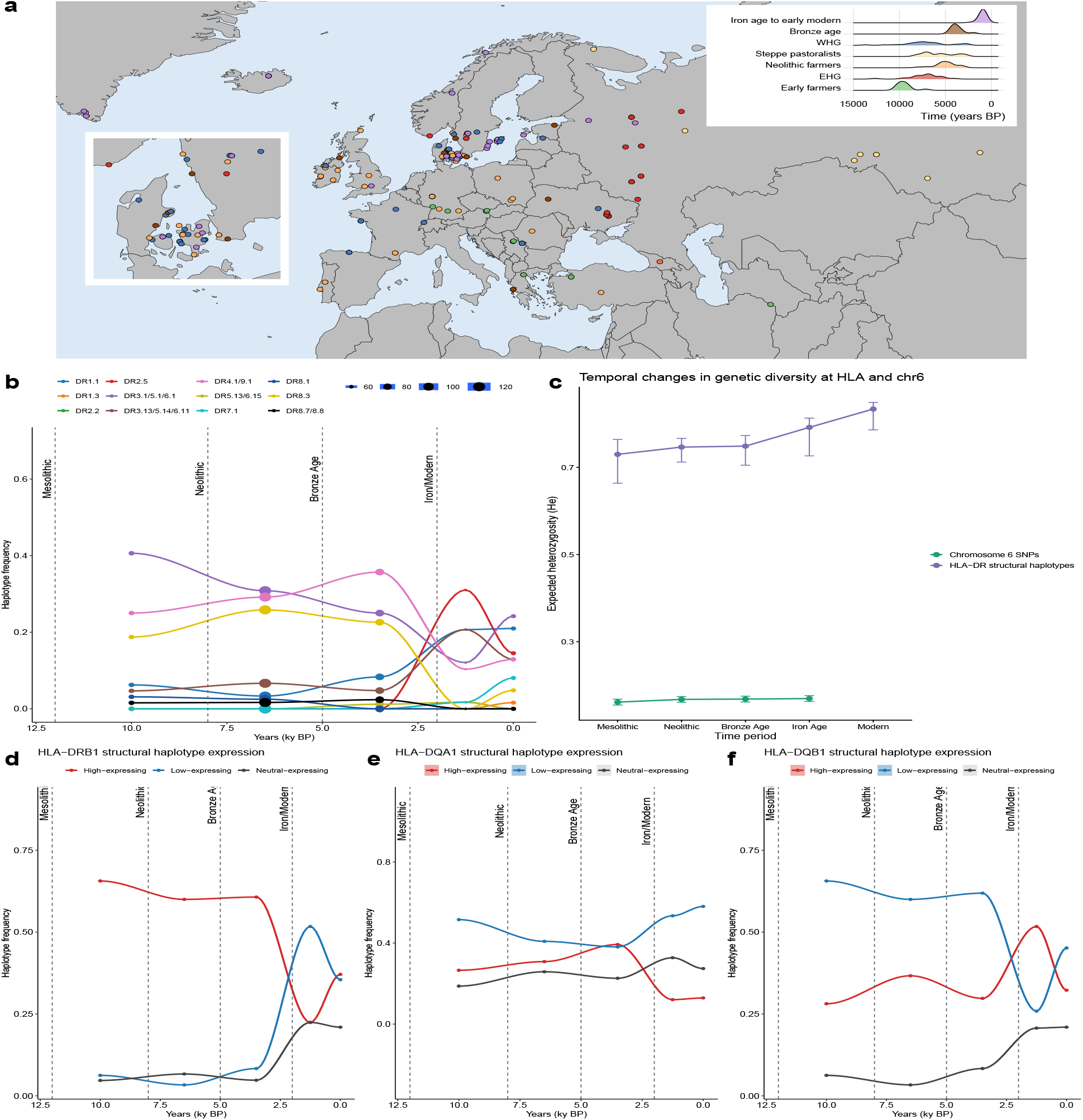
HLA-DR-DQ structural haplotype frequencies in 168 ancient European genomes. **a** Geographic origin, dating and demography of the 168 ancient European genomes included in this study. Each color represents a distinct ancient population. The inset represents the geographic coordinates for Denmark, the country with the most samples. WHG: Western Hunter-Gatherer; EHG: Eastern Hunter-Gatherer. **b** Frequency of HLA-DR-DQ structural haplotypes over time. Time bins (unit in kyr BP): [12,8), [8, 5), [5, 2), [2, 0.5), [0.5, 0]. The points represent the average within each bin. Dot size for each bin represents the sample size. Each bin represents a different epoch (Mesolithic, Neolithic, Bronze age, Iron age and Modern age, respectively). **c** Expected heterozygosity (He) of HLA-DR-DQ structural haplotypes and SNPs from 100 200 kb non-HLA regions on chromosome 6 across archaeological periods. Points represent mean He per period, with error bars denoting 95% bootstrap confidence intervals for HLA-DR-DQ structural alleles while denoting 95% confidence intervals for chromosome 6 SNP allelic diversity from the 100 distinct analysed regions. Modern SNP analysis is not included as the VCF file from ref. (*79*) does not include modern genomes. HLA-DR-DQ diversity increases more strongly in later periods relative to background genetic diversity which sees a slight increase in earlier periods. Structural haplotypes were also grouped into *DRB1* (**d**), *DQA1* (**e**) and *DQB1* (**f**) modified gene expression (table S7) and the frequency shifts over time as done in **b**. *DRB1* high-expressing and *DQB1* low-expressing trends share similarities as both DR4.1/9.1 and DR3.1/5.1/6.1, the most common haplotypes in ancient genomes, have the same directional effect on gene expression across these genes.

By grouping samples into broad archaeological periods, we found that DR-DQ structural haplotypes DR4.1/9.1, DR3.1/5.1/6.1 and DR8.3 were the most prominent from the Mesolithic through to the Bronze Age and subsequently significantly declined in frequency (linear regression, *p*-value = 8e-6, 5.9e-7 and 1.2e-7 for DR4.1/9.1, DR3.1/5.1/6.1 and DR8.3, respectively; Fig.5b). This decline, we noticed, came with an increase in allele diversity (figure S23-24). Expected heterozygosity of HLA-DR-DQ structural haplotypes increased progressively from the Mesolithic to recent historical periods (Spearman’s *ρ, p* = 0.016; Fig.5c), with early populations dominated by a smaller number of haplotypes and later populations exhibiting a more even frequency distribution, largely driven by a marked increase in diversity from the Iron Age onward. This trend was not driven by unequal sample sizes across periods: earlier periods included more individuals than later ones.

To assess whether this pattern could be explained by demographic processes alone, we compared HLA-DR-DQ structural diversity trends to those observed in 100 putatively neutral 200 kb regions on chromosome 6 outside the extended HLA region. While diversity in these background regions also increased over time, particularly from the Mesolithic to the Neolithic, the temporal trajectory differed significantly from that of HLA structural haplotypes. Specifically, a slope comparison excluding Modern frequencies revealed significant differences (*p* = 0.031; Fig.5c), indicating non-parallel trends and suggesting that demographic expansion alone cannot fully account for the observed increase in HLA-DR-DQ structural haplotype diversity.

To explore temporal patterns in HLA gene expression, we classified structural haplotypes according to their association with gene expression levels and tracked their frequencies through time. Haplotypes were grouped as high-, low-, or neutral-expression based on significant differences in expression (Dunn’s test with Benjamini–Hochberg correction; table S7; figure S25–27). Haplotypes associated with high *HLA-DRB1* expression were prevalent from the Mesolithic through to the Bronze Age but declined during the Iron Age (Fig.5d), coinciding with the period in which overall structural haplotype diversity increased. In contrast, for *HLA-DQA1*, haplotypes associated with high and low expression showed largely overlapping temporal trajectories, and no clear directional trends were observed (Fig.5e). Finally, for *HLA-DQB1* we saw a decrease in lower-expressing haplotypes and an increase in neutral-expressing ones (Fig.5f).

## 3 Discussion

The HLA region is one of the most polymorphic regions in the human genome. This diversity is thought to be maintained by pathogen-mediated selection, as variation in HLA genes influences antigen presentation and downstream immune activation. While the SNPs and indels across the region have been well characterised, the structural variation found across the region has remained an open area of genomic research. With complete HLA haplotype sequences, we have been able to better uncover the extent and complexity within this region (*5, 13, 33*). Our study provides a comprehensive understanding of the structural diversity across the HLA locus through an analysis of 460 haplotypes. This revealed that human HLA haplotypes have an average of 68 SVs which can span multiple HLA genes and pseudogenes. However, we find most structural complexity to accumulate across the HLA-DR-DQ region, which is in strong linkage disequilibrium. Our annotations also reveal novel HLA alleles predominantly found in non-European populations. These results highlight the need to increase the discovery of novel alleles and structural variants in under-represented populations. Despite the diversity we uncover, most structural variation is globally shared rather than population-specific, consistent with long-term balancing selection maintaining haplotype diversity. The use of a pangenome reference graph enabled improved representation of large SVs. As minigraph requires a backbone genome, some degree of reference bias may persist; however, the use of the complete T2T CHM13 assembly likely mitigates this effect.

The resolution of 13 structural haplotypes across HLA-DR-DQ confirms that *DRB1* intronic and exonic remnants are major contributors to structural heterogeneity. We show that these structural haplotypes are significantly associated with variation in HLA class II gene expression, establishing SVs as an important, historically overlooked, factor in HLA regulation. We then go on to look for specific SVs that are driving this differential gene expression, revealing 168 SVs associated with modified classical HLA genes’ expression. The addition of SVs to SNP-based models increased the explained variance in gene expression, particularly for class II HLA genes, underscoring the important regulatory contribution of SVs in these loci. Since *DRB1* gene expression has been established as an important factor determining vaccine response (*9*) and infectious disease severity (*34*), our findings suggest that variation in gene regulation driven by SVs may have functional consequences for immune phenotypes. Accordingly, future GWAS, autoimmune and infectious disease studies, and cancer research should consider the contribution of SVs to gene expression and immune function. More broadly, disease susceptibility may arise not only from variation in peptide-binding residues but also from structural effects on gene regulation. As high-quality phased assemblies continue to expand through efforts such as HPRC, we anticipate further resolution of the mechanisms linking HLA structure, regulation, and immune phenotypes. We note, however, that our expression analyses rely on Iso-Seq and RNA-Seq data generated from B-lymphoblastoid cell lines (BLCLs). Expression patterns measured in BLCLs may not reflect in vivo regulation or cell-type–specific responses. Allele-specific expression at HLA class II loci also changes during T-cell activation (*8*), meaning our results capture baseline transcriptional effects of SVs but not the full regulatory dynamics and cell-type specificity during immune stimulation. Epigenomic influences likely contribute additional layers of regulatory variation and represent an important direction for future work. Finally, we note that our analyses do not establish causality for the identified SV-eQTLs. Given the extensive linkage disequilibrium and structural complexity of the HLA locus, the associated SVs may tag other regulatory variants, and functional validation will be required to pinpoint causal elements.

We further introduce a pangenomic framework for long-read genotyping of large SVs across the HLA region (Supplementary Text 3; figure S28). This approach increases the sensitivity of insertion detection while maintaining high precision, thereby improving representation of complex insertions in long-read sequencing studies. While Sniffles (*35*) performs robustly for smaller SVs, our results indicate that pangenomic approaches improve detection of large and highly polymorphic structural events that are common in the HLA locus. A limitation of our approach would be that called SVs are not phased, though this will be tackled in future versions of the pipeline by phasing the long reads.

Comparative analysis of telomere-to-telomere non-human primate genomes reveals additional pseudogene complexity and clarifies the origins of human-specific deletions, particularly those surrounding *HLA-A*. In non-human apes, *MHC-A* is essential for class I-mediated peptide presentation (*36*). Our transcriptomic analyses show that SVs upstream of *HLA-A* influence gene expression levels in humans, suggesting that the extensive structural complexity observed in ape genomes may similarly modulate *MHC-A* expression, with implications for non-human primate immune responses, infectious disease modeling (*36*) and conservation genetics (*37*). These annotations provide a foundation for improved immunogenetic analyses in primates and enhance interpretation of human-specific HLA haplotype evolution.

Our study reveals an increase in structural haplotype diversity at the HLA-DR-DQ locus in ancient Europe that cannot be fully explained by demographic history. The HLA-DR-DQ region plays a critical role in immune defense against infectious diseases, and during the Iron Age and later periods marked by high population densities and frequent pandemics, previously rare alleles, some with lower *DRB1* expression, likely became advantageous under these new selective pressures. This aligns with prior evidence that immune gene adaptation intensified after the Bronze Age, coinciding with increasing pathogen loads (*38*). Notably, the DR2.5 haplotype, characterised by low expression and the presence of *DRB1*\*15:01, which has been associated with susceptibility to medieval leprosy (*39*), rose sharply during the Iron Age, possibly introduced by Steppe pastoralists (*40*), before declining in frequency in modern Europeans. Importantly, higher or lower *HLA-DRB1* expression does not necessarily correlate with antigen presentation efficiency, which may explain recent decreases in some high-expressing haplotypes alongside growing overall diversity driven by pandemics. We acknowledge several limitations of this ancient DNA analysis. Genome coverage varies across time periods, and HLA typing, although manually curated and supported by high Optitype scores, remains imperfect in ancient samples. Moreover, our expression analysis assumes that structural haplotypes had the same expression patterns in modern and ancient genomes. Future studies with more high-quality ancient genomes from diverse populations and explicit evolutionary models will be essential to fully understand how shifting demographic and pathogen landscapes have shaped HLA diversity worldwide. Achieving this breadth remains challenging, however, due to the scarcity of well-preserved archaeological remains and recoverable ancient DNA from warmer and more humid regions, which are underrepresented despite their importance for global HLA diversity.

In conclusion, our study provides a detailed characterisation of structural diversity across the HLA locus and highlights the role of SVs in modulating HLA gene expression. Expanding phased assemblies across globally diverse populations will further illuminate the evolutionary, regulatory, and disease-relevant consequences of structural haplotype variation, with broad relevance to GWAS interpretation, autoimmune disease research, and cancer immunology.

## 4 Materials and Methods

### 4.1 Datasets

464 high-quality long-read assemblies were compiled from the HPRC (*5*) intermediate Spring 2025 release as well as the reference genomes CHM13 (*41*) and GRCh38. Haplotype assemblies would only be included if they contiguously spanned the HLA locus. Gaps in the HLA regions were detected by aligning the assemblies to the CHM13 reference (see HLA extraction and annotation) as well as by looking at the assembly quality control procured by the HPRC. After excluding 4 assemblies that presented gaps or misassemblies across the locus, the final number of analysed assemblies would be 460 as well as the reference genomes GRCh38 and CHM13 for certain analyses.

The Iso-Seq data also came from the HPRC Release 2, which contained flnc.bam files (PacBio RNA-Seq data with removed primers and polyA tails) for 199 individuals. The short-read RNA-Seq data was taken from 27 samples that were also found in the Geuvadis project (*28*). 50 further RNA-Seq samples were obtained from multiple projects (*6, 29, 30*), bringing the total available RNA-Seq datasets to 77.

We used ONT long-read datasets from 976 individuals to further characterise the structural variation found across the HLA region in diverse human populations (*20*). We used a subset of 42 samples from that dataset that coincided with the HPRC phase II samples for the long-read SV calling benchmark.

### 4.2 HLA extraction and annotation

HLA regions were extracted from each assembly by aligning each respective chromosome 6 using wfmash (*42*) (v0.22.0) in an all-to-CHM13 alignment and extracting the HLA coordinates from each chromosome using impg (*26*) (v0.2.3) “impg query -p mappings chr6.paf -r “CHM13#0#chr6:28385000-33500000” -x”. The region selected was surrounding the HLA locus with flanking sequence from the CHM13 genome. RepeatMasker (*43*) (v4.1.7-p1) and Dfam (v3.8) were used to find repetitive elements across each HLA locus. Immuannot (*14*) was used to annotate HLA genes across each HLA locus, meanwhile MHC-annotation (*44*) (v0.1) was used to capture the rest of the gene content (non-HLA genes/pseudogenes). Bedtools (*45*) (v2.30.0) was used to intersect annotation files with themselves to check for wrongly overlapping genes.

### 4.3 Variant calling in the HLA region

Variants (both SNVs and SVs) were called using the Phased Assembly Variant caller (*46*) (PAV; v2.4.6), a leading phased assembly-based SV caller (*13*). As not all large variation was being captured by PAV, we additionally called SVs by creating an HLA-Pangenome reference graph (HLA-PRG) using minigraph (*47*) (see ‘HLA pangenome graphs’ for more details) with SVs ≥ 2,500 bp to only capture large rearrangements in the graph. After manually confirming that large SVs were being captured by the graph (such as the *HLA-HTKU* deletion), each HLA haplotype was mapped back to the HLA-PRG and SVs called using miniwalk (v1.0.1) (*48*). The two VCF files were then merged using Jasmine (*49*) (v1.1.5), keeping unique SVs from both files. VCF files were filtered for variants ≥50 bp and merged across haplotypes using Jasmine with default parameters.

SVs from our dataset were compared against HGSVC3 (*13*) HLA SVs by comparing the SVs across the same range in chr6 (28,385,000-33,500,000) with truvari (*50*) (v4.2.2). We used a reciprocal threshold of 0.7, maximum reference distance of 1,000 bp and sequence similarity of at least 0.5, to allow for more flexibility for minigraph-called SVs. The false positive and false negative VCFs were then further collapsed using truvari (*50*) (v5.4.0) with a maximum reference distance of 10,000 bp and sequence similarity of at least 0.5.

### 4.4 Linkage disequilibrium and recombination rate analysis

To calculate the linkage disequilibrium across the HLA locus in our samples, we initially used bcftools (*51*) (v1.15.1) to filter out SVs and indels from the PAV VCF files and posteriorly merge them. Only biallelic sites were kept (‘-m2 -M2’). To calculate pairwise linkage disequilibrium, we ran plink (*52*) (v1.9b 6.21-x86 64) with options –indep-pairwise 50 5 0.2 to prune low R2 values, to reduce the SNP density. We then reran plink using –r2 square –allow-extra-chr —maf 0.05 –from-bp 3880000 –to-bp 4600000 –chr CHM13#0#chr6, for capturing LD across the HLA genes in class II. This process was done for the global VCF file as well as for the population-specific VCF files. The resulting genotype matrix with the physical positions of SNPs for linkage disequilibrium calculation (R^2^ statistic) was plotted with the LDheatmap (*53*) function in R (v4.4.3).

To calculate the recombination rate across the HLA region we downloaded the genetic recombination map by ref. (*54*) and used liftOver (*55*) to change the coordinates from GRCh38 to CHM13 as the SVs were genotyped against CHM13.

### 4.5 HLA pangenome reference graphs

As mentioned in “Variant calling in the HLA”, an initial HLA-PRG was created to aid in calling large SVs, possibly missed by PAV. This graph was created using minigraph (v0.21) with the -cxggs-l 2,500 flags to only include large SVs. The backbone of the HLA-PRG was the reference genome CHM13 and the rest of 461 haplotypes were added to the PRG in lexicographic order, starting with GRCh38. We created a second PRG (HLA-PRG2) with the -l 50 flag modified to include all types of structural variation. All haplotypes were mapped back to HLA-PRG2 using minigraph to determine the paths traversed by each haplotype across the graph in the HLA class II region for structurally variable region identification across our dataset while class I was determined by mapping the haplotypes to HLA-PRG.

### 4.6 Haplotype clustering and annotation

To extract the haplotype sequences from the graph (HLA-PRG2), we concatenated the nodes traversed by each haplotype to create a fasta sequence using a custom script. We then wanted to determine how similar the haplotypes were both within and between DR groups. We used MAFFT (*56*) (v7.505) to do a multiple sequence alignment for the haplotype sequence within each DR group. We then used distmat (*57*) (EMBOSS v6.6.0.0) to obtain a distance matrix for each DR group. The clustering threshold of 0.03 was decided after visual inspection of the alignments. MAFFT was also used to align all final 20 haplotypes. To construct a phylogeny of the main haplotypes, we used IQtree (*58*) (v2.0.7) with the general time-reversible model with rate heterogeneity and DR7.1 as the outgroup as it was the largest haplotype and evolutionarily one of the oldest (*12*).

To annotate each haplotype and determine its gene content and lengths, the haplotype traversal paths through the minigraph graph (HLA-PRG2) were extracted and visualised using Bandage (*59*) as well as all known genes and pseudogenes across the regions. Solitary exons/introns were determined by both visualisation using Bandage and by mapping sequences suspected of containing exons/introns to the IMGT/IPD database (*3*) (v3.56.0) holding genomic sequences of the *HLA-DRA*-*HLA-DQB1* genes using BLASTN (*60*) (v2.12.0).

The same pipeline was applied for the structural haplotypes spanning *HLA-A* and *HLA-B*/*HLA-C*, except the haplotypes were determined using HLA-PRG due to the lower structural complexity in those regions.

Transcription factor binding site motif predictions were calculated for each extracted haplotype using MEME Suite’s (v5.5.7) FIMO (*61, 62*). Motifs were obtained from the JASPAR2022 CORE vertebrates non-redundant database. Input sequences in FASTA format were scanned using default parameters with a p-value threshold of 1e-4 to identify statistically significant motif matches.

### 4.7 Ape MHC annotation

To annotate the pseudogenes across the *HLA-A* region in the ape T2T genomes, we used the human genome sequences. We manually annotated the pseudogenes through two parallel methods. First, we extracted the ape MHC regions by mapping the ape T2T assemblies to the CHM13 human genome using wfmash and extracted the MHC using impg. The MHC ape sequences were then added on to HLA-PRG2 with minigraph to study structural rearrangements. We then visualised the ape MHC pangenome and mapped the human HLA gene and pseudogene sequences to it using Bandage (*59*). Second, we used exonerate (*63*) (v2.4.0) with the “affine:local” mode to map the human HLA genes as well as the ape genes annotated in IPD/IMGT (v3.14) across the ape haplotypes. We confirmed our annotations for functional genes using the “protein2dna” mode with IPD/IMGT (v3.14) protein sequences. Pseudogene annotations were confirmed where both methods coincided.

### 4.8 Ape *MHC-A* phylogeny

To create a phylogeny of the 460 sequences and the ape T2T genomes (excluding the siamang genome) (*26*) we used the *HLA-A* sequences as the gene was found in all ape genomes and was flanking the 4 main human structural haplotypes. We constructed the phylogenetic tree using IQtree from a multiple sequence alignment done using MAFFT (default parameters) across all 470 sequences with the general time-reversible model with rate heterogeneity and 1,000 bootstraps.

### 4.9 Gene expression analysis

Iso-Seq flnc.bam files were downloaded, along with each sample’s both phased assemblies for 199 individuals, excluding 3 individuals without both HLA haplotypes, as read phasing wouldn’t have been possible. For the remaining individuals, we mapped and sorted the reads to both of their corresponding assemblies using pbmm2 (v1.17) (*64*). We then filtered the bam files to keep only the HLA region in each assembly using samtools (*51*) (v1.21). The reads were phased using a custom Python script, where each read mapped across the HLA region was compared to its respective mapping quality in both assemblies using the CIGAR string, a method already proven to be accurate when phasing Iso-Seq reads (*13*). Matches across the CIGAR string were evaluated positively, “Ns” neutrally while mismatches, deletions, or any other attribute was evaluated negatively across the overall alignment score. Whenever a read scored better in one haplotype over another, we associated it to the higher-scoring one, meanwhile exact scores would imply reads that could not be phased. Reads that were uniquely mapped to an assembly were phased for that assembly. A list of reads was extracted from this analysis to phase the mapped bam files to keep only the relevant reads for each assembly. The filtered, phased, bam files were converted to fastq to be input to IsoQuant (*65*) where the CHM13 genome was used as reference as well as its HLA gene annotations. IsoQuant output gene and transcript quantification in the normalised transcripts per million (TPM) format. This output was used to evaluate mono-allelic gene expression across samples.

To evaluate the role of structural variation affecting gene expression, for each sample we created a personalised CHM13 reference genome which included the HLA gene sequences of each sample’s haplotypes while masking CHM13’s HLA genes. Reads were mapped to the personalised reference genome using pbmm2 (“ISOSEQ” preset) with the resulting bam file being input to featureCounts (*66*) for gene expression quantification using an enhanced annotation file which included the additional personalised HLA annotations. A minimum mapping quality of 20 was used as a filter. Those samples that presented close to perfect phasing (<5% unphased reads) were used to determine haplotype-specific HLA gene expression. For the short-read RNA-Seq analysis, we trimmed and filtered low-quality reads using fastp (*67*) (v0.23.2), mapped to the personalised genome using STAR (*68*) (v2.7.10) and calculated gene quantification with featureCounts, with the paired-end mode. We calculated transcripts per million (TPM) to normalise the expression values across samples. The length for each haplotype’s HLA transcript to calculate the TPM was inferred from each haplotype’s CDS allele length taken from the IMGT/HLA database. We adopted this approach as previous pipelines establishing personalised references for HLA gene expression quantification and masking reference HLA sequences and transcripts have been successful (*9, 31*). Moreover, our HLA TPM values were in line with what has been previously observed (*31*).

We detected cis SV-eQTLs across the HLA region that were associated with HLA gene expression for the 7 classical HLA genes that could be phased (*-A,-B,-C,-DRB1,-DQA1,-DQB1* and *-DPB1*). Tests were done independently per gene as, due to phasing, differing number of haplotypes were available for each gene. Covariates used for the analysis included the first three principal components calculated from the SNP data of the 77 individuals (1KGP phase III) (*69*), 15 probabilistic estimation of expression residuals (PEER) (*70*), sex and batch. Covariates were duplicated for each of the phased samples which led us to test associations using a mixed model to account for the non-independence of the phased samples. We applied the Benjamin-Hochberg correction (*p*-value < 0.05) to the resulting *p*-values. This methodology was repeated for the SNPs. Only variants with MAF > 0.05 were included in the association analysis.

To quantify the change in explained variance by accounting for SVs, for each gene, we first fit a baseline model including the five most strongly associated SNPs identified from single-variant association analyses. We then fit a nested model that additionally included the five most strongly associated SVs for the same gene. The proportion of variance explained by fixed effects was quantified using marginal R^2^, and the incremental contribution of SVs was assessed as the change in marginal R^2^ between SNP-only and SNP+SV models.

### 4.10 HLA typing benchmark on modern and ancient short-read sequencing data

The genotyping performance for Illumina data was evaluated for HLA genes’ alleles. We included individuals whose both HLA haplotypes were complete, making for a total of 227 individuals with phased genotypes included as a reference. Locityper (*71*) (v0.19.1) was used for HLA typing. The locityper database was created on 20 HLA genes and their respective coordinates in CHM13: *HLA-A, HLA-B, HLA-C, MICA, MICB, HLA-DRA, HLA-DRB1, HLA-DQA1, HLA-DQB1, HLA-DPA1, HLA-DPB1, HLA-DMA, HLA-DMB, HLA-DOA, HLA-DOB, HLA-DQA2, HLA-DQB2, HLA-DPA2, HLA-DPB2, HLA-G. HLA-DRB1* and *MICA*’s coordinates were expanded to accommodate variants going beyond the gene coordinates. We used a leave-one-out approach to avoid bias towards test samples already being present in the graph. SNVs, indels and SVs were recovered from the PAV VCF file for each sample, across those genes, where individual haploid VCFs were merged with their corresponding haplotype into diploid VCFs before merging with the rest of samples using bcftools (v1.15.1) and bedtools (v2.30). Genotyping performance was evaluated on the 4 different fields of the HLA nomenclature. Filtering of *HLA-DRB1* calls was done by keeping individuals with quality calls > 21 (median value for exact matches) and weight distance < 0.04 (median value for exact matches; Extended Data Fig.6c).

We used a subset of 40 modern genomes presenting most HLA structural groups to simulate ancient genome sequence profiles, to determine whether Locityper’s genotyping approach is appropriate for low-coverage, error-prone samples. A similar approach as in ref. (*72*) was taken. NGSNGS (*73*) was used to simulate the ancient genome single-end reads using the following parameters: `-f fq -s 4 -ne -lf Size dist sampling.txt -seq SE -m Illumina`, `0.024, 0.36, 0.68, 0.0097 -q1 AccFreqL150R1.txt`, with varying coverages ranging 1-10X. The reads were created from each sample’s respective chr6 haplotypes as well as CHM13’s chr17, as Locityper relies on mapping reads in a simple region in chr17 without SVs, for read coverage normalisation (*71*). Simulated reads shorter than 70 bp were filtered out as strobealign (*74*), Locityper’s short-read aligner, does not handle unusually short reads well. We ran the simulated data through both Locityper and OptiType (*32*). For the ancient HLA typing benchmark we did not apply a leave-one-out approach as it was not necessary; we were focusing on typing the first field of specific HLA genes which was not unique to any one sample. As we were mainly interested in the 13 main structural haplotypes that could be inferred from the first field of *HLA-DRB1* and *HLA-DQA1* typing, we focused on comparing the first field between expected and observed typing results. OptiType mainly types *HLA* class I genes, however, a development version of the tool is currently able to also type class II genes. We modified the allowed parameters of the tool to also allow single-end reads, which is the case for most ancient DNA.

All benchmarking has been wrapped in a Snakemake pipeline for reproducibility.

### 4.11 HLA structural variant calling from long-read pangenome alignments and benchmarking

Long-read sequencing data (ONT) from ref. (*20*) were aligned to the reference CHM13 using minimap2 and reads mapping across the HLA region were extracted using SAMtools (v1.21). Those reads were then mapped to the HLA-PRG using minigraph. To distinguish heterozygous from homozygous variants, we developed a depth-based genotyping pipeline that accounts for non-uniform coverage across the graph. For each sample, reads mapping to graph nodes were counted, filtering alignments where neither the read nor node achieved ≥90% coverage to exclude spurious mappings. Multi-node alignments were decomposed such that each traversed node received one read count. We calculated per-node sequencing depth by partitioning nodes into 100 bp bins (or base-by-base for nodes <5kb) and computing the mean coverage across positions, accounting for the precise mapping coordinates of each read within the node.

Core nodes, those flanking all bubbles in the pangenome graph, were identified from the gfatools (*75*) (v0.5-r292-dirty) bubble output, and their average depth (for nodes ¿150 bp) established the expected homozygous coverage with 95% confidence intervals. For alternate allele nodes within each bubble, depth ratios relative to flanking core nodes were used to classify variants: nodes with depth >0.6× core coverage were called homozygous (present on both haplotypes), 0.3-0.6× heterozygous (one haplotype), and <0.3× absent. Haplotype paths were constructed by separating homozygous nodes (assigned to both haplotypes) from heterozygous nodes, which were distributed between haplotypes based on graph topology. Specifically, heterozygous nodes were assigned to haplotypes by counting available (non-occupied) edges to existing paths in each haplotype. Final node ordering within each haplotype respected the directed edges in the GFA file, with topological sorting of connected components and numerical ordering as a tiebreaker for disconnected nodes. This approach generated unphased BED files representing the genomic paths traversed by each haplotype through the pangenome graph. This pipeline has been established as a new miniwalk module.

For calling SVs, the same pipeline as established previously in “Variant calling in the HLA region” was used to obtain two VCF files for each unphased haplotype. The haplotypes were merged using SURVIVOR (*76*) (v1.0.7) and the VCF was further refined using a custom Python script. The final VCF was benchmarked against the SVs found in “Variant calling in the HLA region” for each of the 41 samples used for the benchmarking using truvari with the parameters “-r 1000 -p 0 -P 0.3-O 0.25 -s 2500 -S 2500 –sizemax 150000 –no-ref a”. SVs across the HLA were also called using Sniffles (*35*) (v2.7.1) to compare the precision and recall against our pangenomic approach. We repeated the benchmark using an HLA graph with large structural variation without the 41 samples used in the benchmark, therefore including 378 haplotypes in the graph.

All the SV benchmarking has been wrapped as a Snakemake pipeline.

### 4.12 Ancient genome analysis

We downloaded 1,290 ancient genomes across Europe and mapped them to the CHM13 reference genome using bwa-mem2 (*77*). We then kept those genomes that had a read coverage of > 0.5X across the HLA region totaling 556 ancient genomes. We typed the HLA class I and class II genes using Optitype as described in “HLA typing benchmark on modern and ancient short-read sequencing data.” We retained samples only if the typing results were consistent; for example, when the *DRB1* allele matched the expected presence or absence of *DRB3, DRB4*, or *DRB5* alleles, and all classical HLA genes typed by Optitype were present. We kept ancient samples that had coverage across the HLA region higher than 2X. This threshold was determined from previous literature (*78*) as well as the heterozigosity rates by comparing with the simulated dataset’s coverages (figure S22). Moreover, we manually verified that the observed typing schemes were consistent with expectations based on linkage disequilibrium patterns from modern genomes; for example, all DR4.1/9.1 structural haplotypes were expected to be accompanied by DQA1*03. After filtering, we had 168 ancient genomes (*79–101*) (table S6). As all our samples from Iron age to early modern came from Northern Europe, as quality control, we calculated the haplotype trends over time uniquely in that region and found similar trends when including all samples, indicating that sampling distribution likely wouldn’t bias our results (figure S29).

For each time period, we calculated expected heterozygosity (He) of HLA-DR-DQ structural haplotypes as 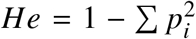, where pi denotes the frequency of the i-th haplotype. To account for uneven sampling and the non-independence of haplotypes within individuals, we estimated confidence intervals for He using a bootstrap procedure that resampled individuals with replacement within each time period (1,000 replicates). In each replicate, all haplotypes belonging to resampled individuals were retained and He was recalculated. Temporal trends in haplotype diversity were assessed by comparing He across time periods and by testing for monotonic trends using Spearman’s rank correlation. Chromosome 6 expected heterozygosity was estimated from autosomal SNP genotypes across 100 regions of 200 kb (approximately the same size as the HLA-DR-DQ region) using published variant calls for a subset (62%) of the same ancient individuals (*79*). For each SNP, allele frequency was calculated within each temporal bin, and per-locus expected heterozygosity was computed as 2*p*(1 − *p*). The 100 regions did not overlap low SNP density regions which could bias the mean He (range 1,230 - 3,368 SNPs per 200 kb region). Chromosome 6-wide He was obtained by averaging across loci within each time bin. To test whether temporal trends in diversity differed between HLA structural haplotypes and the genomic background, we fitted a linear model of expected heterozygosity as a function of time, data source (HLA-DR-DQ vs chr6-wide), and their interaction. Time was treated as a continuous variable using time bin midpoints. A significant interaction term was interpreted as evidence for non-parallel temporal trends.

To determine whether the structural haplotypes’ effect on gene expression could be seen over time across the ancient genomes, we grouped them into high, low or neutrally HLA gene-expressing, dependent on the gene being studied (table S7).

### 4.13 Statistics and visualisations

Unless stated otherwise, all statistics and visualisations were done in R (v4.5.1).

## Acknowledgments

We thank José Antonio Barraza López for assistance with figure design and image preparation. We also thank Hardip R. Patel for relevant discussions regarding the manuscript.

## Author contributions

A.C.-B. conceived this project with input from M.S., P.M.S., L.J.M.C. and S.J.D. A.C.-B. ran all the bioinformatics experiments and analyses with assistance from S.F.J. The work was supervised by M.S., P.M.S., L.J.M.C. and S.J.D. A.C.-B. wrote the first draft which was revised by all authors.

## Data and materials availability

All PRGs generated for this study (HLA-PRG, HLA-PRG2 and Ape-MHC-PRGs), the *HLA-A* ape phylogenetic tree and the 460 HPRCv2 samples’ SV VCF file is available at https://doi.org/10.5281/zenodo.18397440. Miniwalk can be accessed through https://github.com/aleixcanalda/miniwalk and the code used for this manuscript can be found in https://github.com/aleixcanalda/HLA_PRG_Paper.

