## Supplementary Information for "Structural variation shapes regulatory and evolutionary diversity at the HLA locus"

**This PDF file includes:**

Materials and Methods

Supplementary Text

Figures S1 to S32

Tables S1 to S7

### Materials and Methods

#### 1. HLA SV calling with Pacbio Hifi long-read datasets and recall calculation

Pacbio Hifi DeepConsensus long reads for each sample were mapped to CHM13 with pbmm2 (*64*) and subset the alignments to the HLA region using SAMtools (*51*) (v1.21). We then called SVs with Sniffles (v2.7.1), DeBreak (v1.0.2; parameters "–rescue_large_ins –poa") and PBSV (v2.11.0). Each VCF, for each of the 15 samples, was then benchmarked using truvari with the parameters "-r 1000 -p 0 -P 0.3 -O 0.25 -s 50 -S 50 –sizemax 150000 –no-ref a".

### Supplementary Text

#### 1. Validation of SV calls using long-read Pacbio Hifi datasets

We wanted to further validate a subset of our SV calls using multiple algorithms based on Pacbio Hifi long-read datasets used to generate the phased assemblies. We found that 66% - 87% (mean = 0.75), 10% - 67% (mean = 0.43) and 65% - 84% (mean = 0.74) of SVs could be validated across 15 individuals for Sniffles (*35*), DeBreak (*102*) and PBSV (*103*), respectively. These results indicate that the majority of SVs identified from our assemblies are independently detectable using long-read data, supporting the accuracy of our SV dataset.

#### 2. Linkage disequilibrium analysis

Linkage disequilibrium (LD) has been a prominent focus of genetic research over the past two decades (*104, 105*), with HLA imputation playing a crucial role in genome-wide association studies (*106, 107*). Although LD across the HLA region has been extensively studied, estimates have largely relied on sparse SNP or microarray panels due to the difficulty of accurate variant calling and phasing in this highly repetitive region using short-read data. To address this, we utilised the fully-resolved haplotypes from the HPRC to identify LD patterns across the HLA and explore which structural haplotypes are likely to be co-inherited. Across HLA class II, we observed that structural haplotypes spanning*HLA-DRA–HLA-DQB1* formed a block with elevated LD relative to the surrounding region, with $R^2$ average pairwise values ranging from 0.09 in Africans to 0.16 in South Asians. This is approximately twice the mean LD observed across the broader class II region (0.047–0.075), indicating that this segment represents a distinct unit of inheritance (Fig.S30; Table.S2). When incorporating recombination rates across the whole HLA region (*54*), we found that the LD block spanning *HLA-DRA-HLA-DQB1* was one of the regions with lowest recombination rates (Fig.1c). For HLA class I, across the *HLA-C-MICB* region we observed the highest recombination rates, dividing the region into two LD blocks, one encompassing *HLA-C-MICA* and another *MICB* (Fig.1c). Across *HLA-A*, however, the recombination rates were lower, indicating the possible presence of conserved haplotype blocks.

**3. HLA genotyping benchmark**

The highly polymorphic and repetitive nature of the HLA region poses significant challenges for short-read-based HLA genotyping (i.e. typing the HLA genes). Nonetheless, pangenomic approaches, such as Locityper (*13, 71*), have shown promise for genotyping complex loci using Illumina data, provided a suitable reference variant set is available. While the largest reference to date, from the HGSVC3 and HPRC Year 1, included 224 haplotypes, the HPRC Release 2 doubled this number to 460 haplotypes. We leveraged this expanded set to benchmark Locityper on *HLA* typing. As genotyping SVs across the HLA locus with short-read data is complex, we determined whether the found structural haplotypes across HLA-DR could be inferred from the *HLA-DRB1* and *-DQA1* first field typing schemes. We found certain haplotypes to have unique first and second *DRB1* field typing matches, though joint *DRB1* and *DQA1* typing can categorise virtually all structural DR haplotypes (excepting DR8.7 from DR8.8; Supplementary Table 5), which could be useful for inferring structural haplotypes when typing using short-read data.

Locityper performed well on modern human samples across most genes for a full typing match. The worst-performing genes were *DRB1*, *DQA1* and *DQB1* (Extended Data Fig.6a). When comparing solely the first typing field, this disparity was only still observed for *DRB1* (Fig.S31). We determined that those genotypes that obtained exact matches had higher quality and lower weight distance calls (Extended Data Fig.6b). This same trend was observed for *DRB1* (Extended Data Fig.6c), where the proportion of exact matches rose from 31% to 75% when filtering by quality and weight distance (see Methods).

We also wanted to determine how accurately Locityper could genotype ancient samples which led us to simulate ancient DNA datasets from 40 modern human genomes with known haplotypes across various coverages (1-10X). As we were interested in understanding the evolution of the HLA-DR structural haplotypes, which can be inferred from the first *HLA-DRB1* and *-DQA1* typing fields, we focused on the typing performance of that first field. Locityper performed poorly across all ancient samples for *HLA-DRB1*, matching both genotypes in a single ancient sample with 10X coverage (Extended Data Fig.6d). This led us to try another *HLA* typing tool, OptiType (*32*), which showed promise for ancient DNA *HLA* class I typing (*78*). We found that genome coverages 4X-10X had exact matches for all samples at the first field while 1X coverages had exact matches in the first field in 79% of the cases, showing the high performance of this specialised tool, even in low coverages (Extended Data Fig.6e). For *HLA-DQA1*, OptiType performed well with exact matches for coverages 4-10X and 73% exact matches in the first field with 1X coverage.

We further hypothesised that a pangenomic approach would likely prove to be beneficial in genotyping large SVs across the HLA compared to using a linear reference as most long reads cannot span the largest structural variation we detected. We developed an approach using miniwalk (*48*) to genotype SVs using mapped, long reads to an HLA minigraph graph with large (≥2,500bp) variation (see Methods). For this benchmark we used a pangenome with all the samples (460) and another leaving out the 41 samples included in the benchmark (378), to avoid bias, and compared our results against using the linear reference CHM13 and Sniffles2 (*35*). We found that, while our approach was not significantly more precise (0.74 vs 0.76 for Sniffles), it had significantly better recall (0.5 vs 0.44 for Sniffles; two-sided t-test, $p$-value = 0.03; Extended Data Fig.7a). This was likely driven by Sniffles being unable to genotype SVs larger than 30kbp. The low precision (34.5%) for large SVs genotyped by miniwalk was driven by a single false positive SV that was incorrectly called in 19 samples. This sequence is largely flooded with repetitive sequences that span thousands of bps, likely driving spurious mappings (Fig.S32). Removing this SV resulted in 100% precision for large SVs (≥70kbp), found in the DR4, DR9 and DR7 haplotypes. We observed different trends for deletions and insertions: our pangenomic approach had high precision for deletions (0.92 vs 0.76 for Sniffles; Extended Data Fig.7b-d) but lower for insertions (0.71 vs 0.86 for Sniffles). Conversely, we saw low recall for deletions (0.25 vs 0.37 for Sniffles; Extended Data Fig.7b-d) but high for insertions (0.63 vs 0.46 for Sniffles; Extended Data Fig.7b-d).

These results show the trade-offs of adopting a pangenomic approach for genotyping large SVs using long-read data across a structurally complex region.

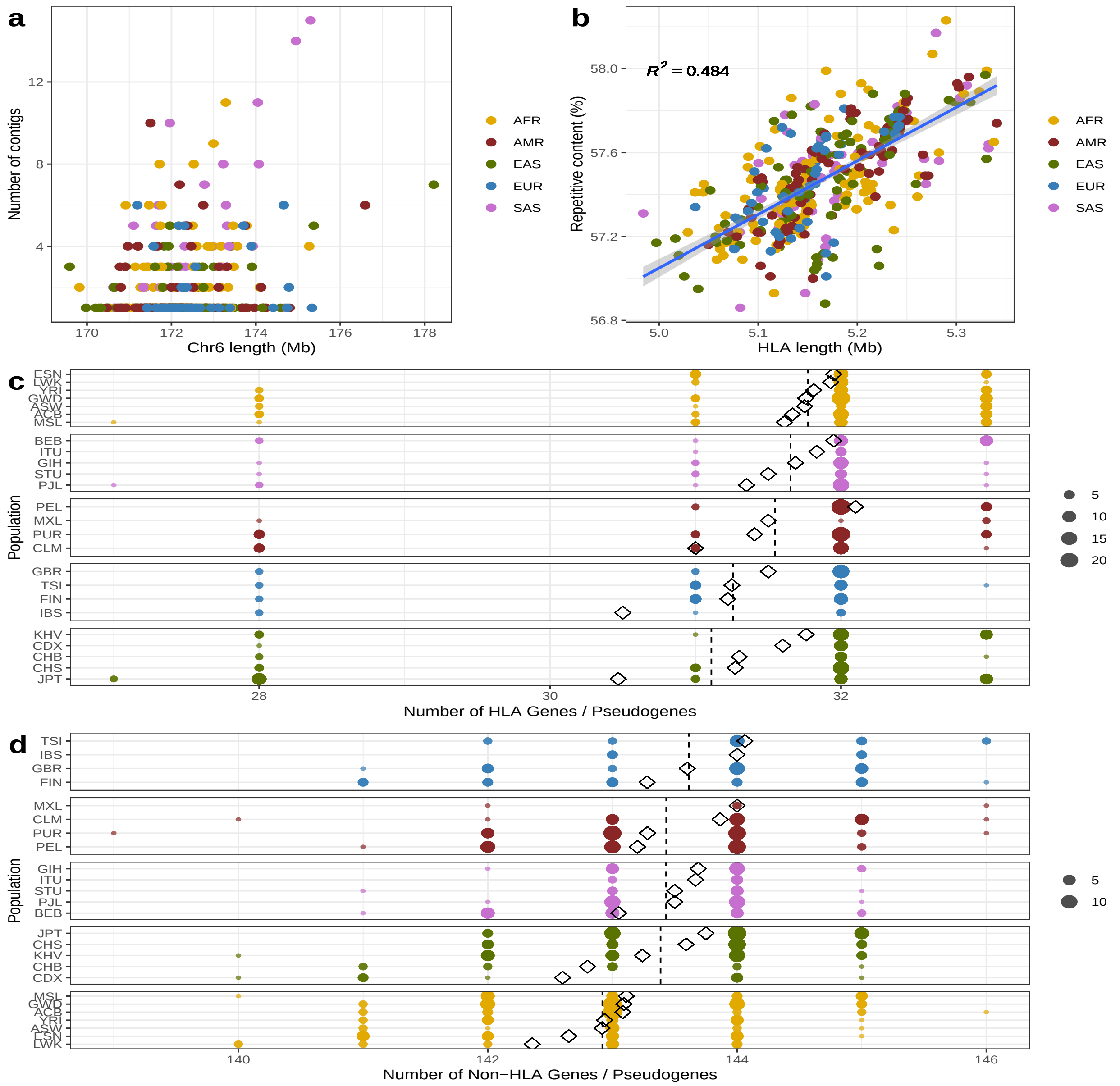


**Figure S1**: **Supplementary Figure 1. Characteristics of 460 complete HLA haplotypes. a** Chromosome 6 number of contigs and length by continental population. **b** HLA length and repetitive content by continental population. A linear model is fit to the data and 95% confidence intervals are indicated by shades. We also analyzed the number of HLA (Immuannot; **c**) and non-HLA (MHC-annotator; **d**) genes, separating by continental populations. Diamonds represent the population means while the vertical, dashed lines represent the continental population means. Continent labels; AFR: African, AMR: American, EAS: East Asian, EUR: European, SAS: South Asian.

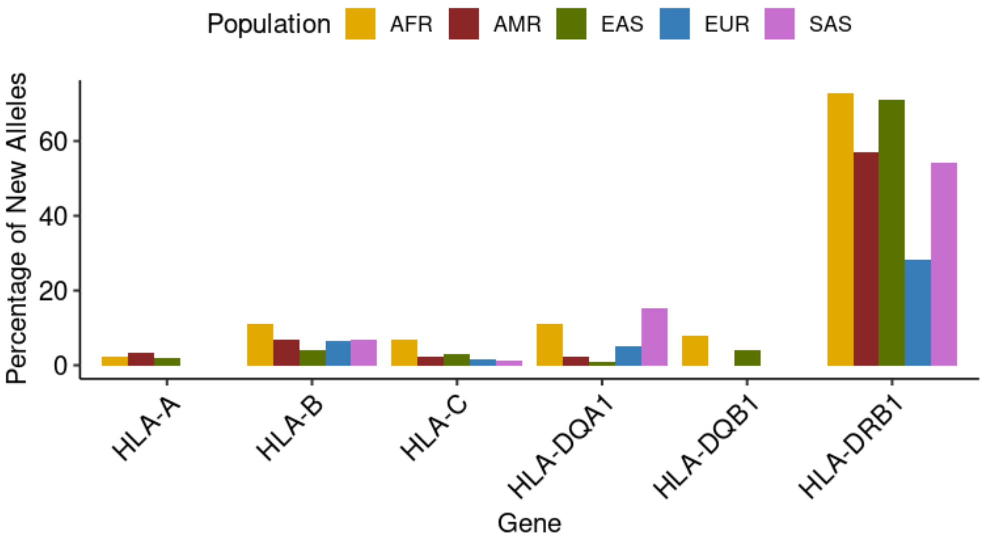


**Figure S2**: **Supplementary Figure 2.** Percentage of newly annotated alleles across the 6 classical class I and class II HLA genes, per continental population. Annotations come from Immuannot.

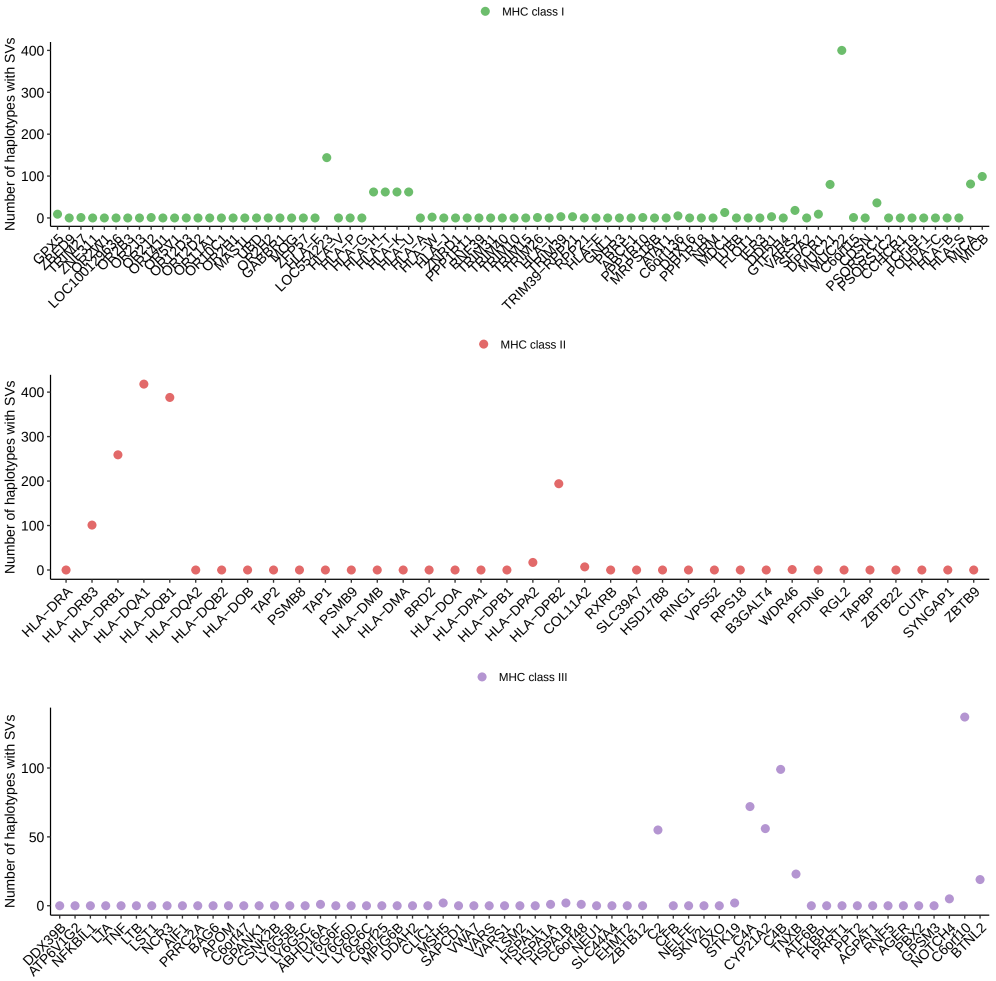


**Figure S3**: **Supplementary Figure 3.** Number of haplotypes with SVs across each HLA gene. Gene annotations done with MHC-annotator.

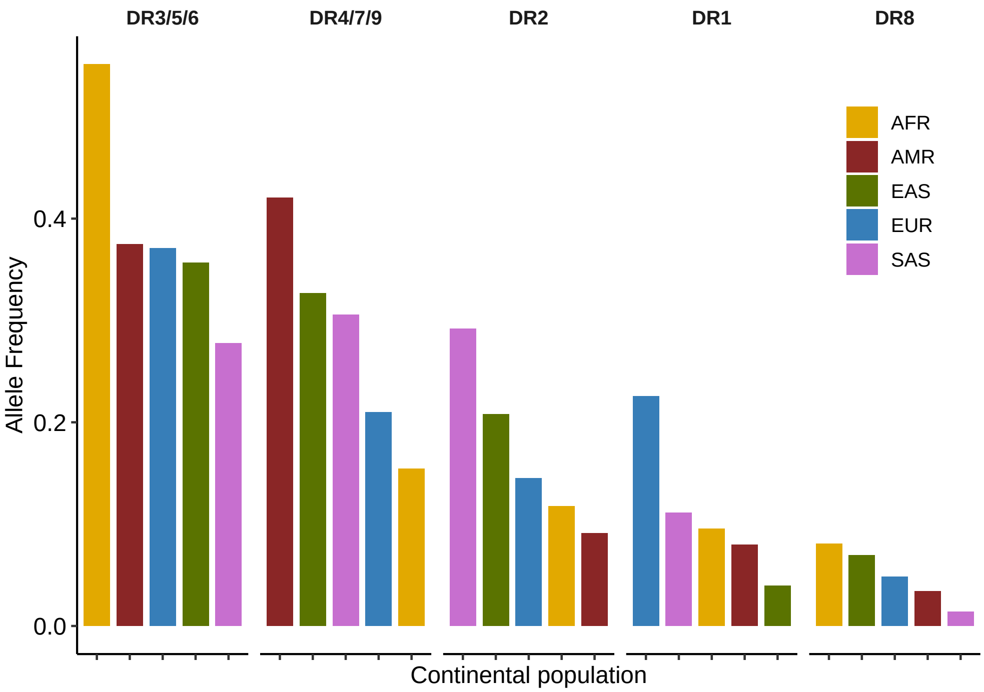


**Figure S4**: **Supplementary Figure 4.** HLA-DR haplotype frequencies across global continental populations.

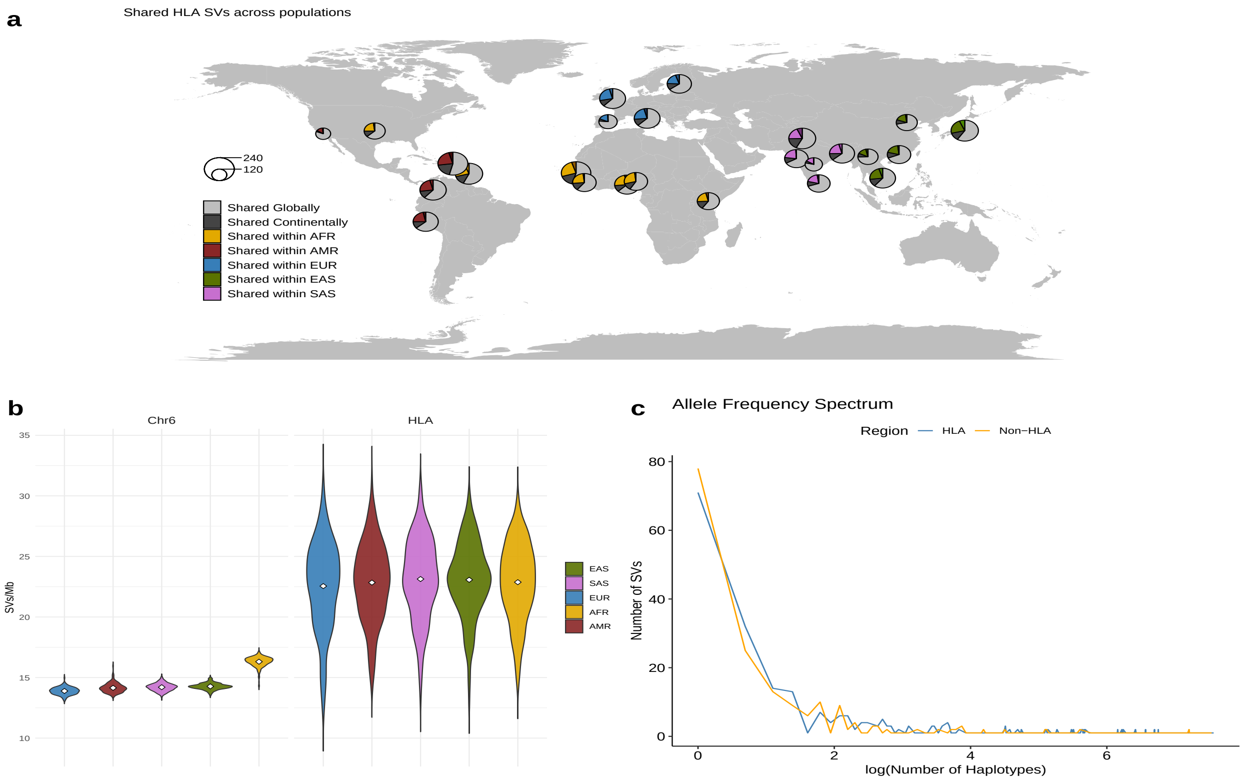


**Figure S5**: **Supplementary Figure 5. HLA SV alleles' global distributions. a** Private and shared HLA structural variation found in phased assemblies across populations. Pie plots are divided into SVs that are found across all populations, those that are found in more than one continent, those that are found within a single continent and those unique to a single population. **b** Violin plots of the SV density (number of SVs per Mb) divided into continental population and genome region (HLA and chr6). White dots represent the mean. **c** Allele frequency spectrum of SVs in the HLA region and out of it in a region of similar size. X-axis represents the log number of samples that share a specific SV, where 1 indicates an SV singleton.

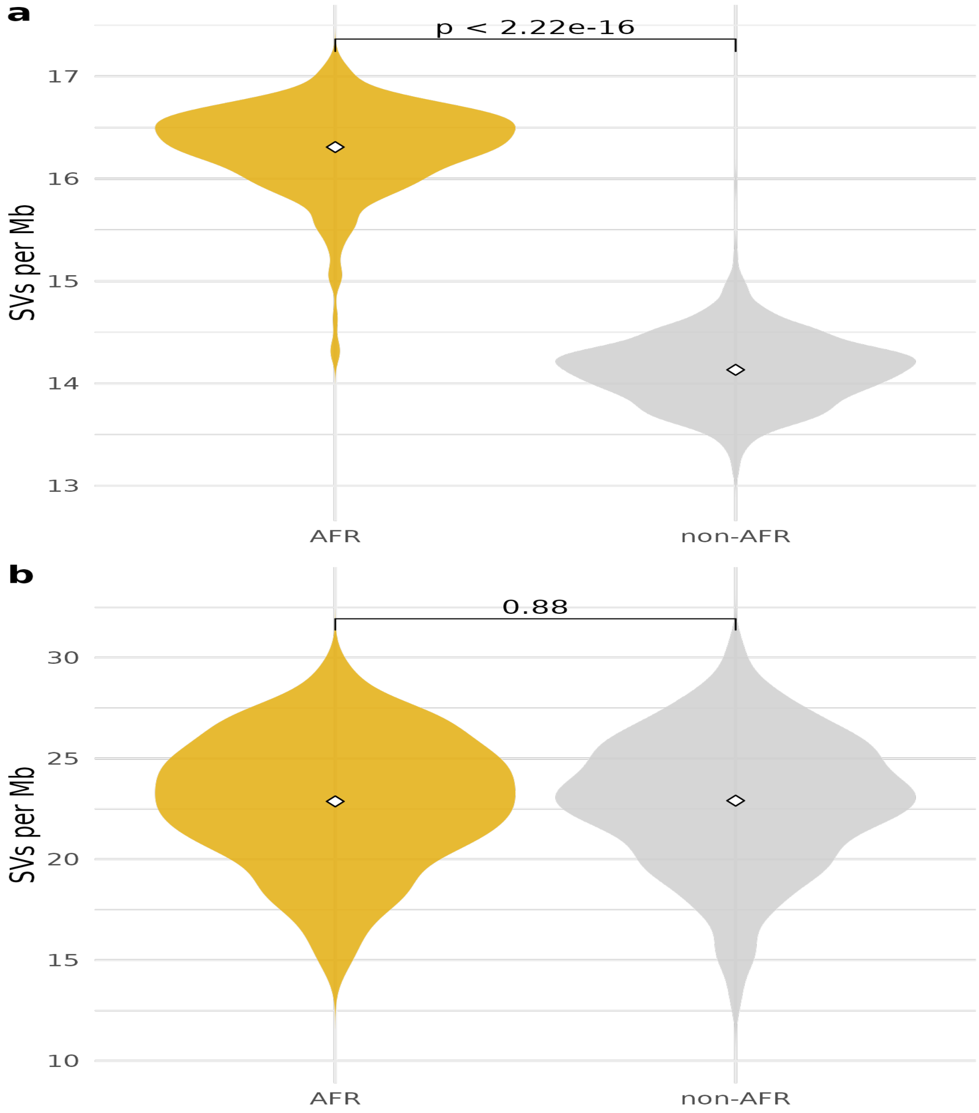


**Figure S6**: **Supplementary Figure 6.** SV density (SVs/Mb) across chr6 (**a**) and HLA (**b**) in African vs non-African populations as violin plots (white dot is the mean value).

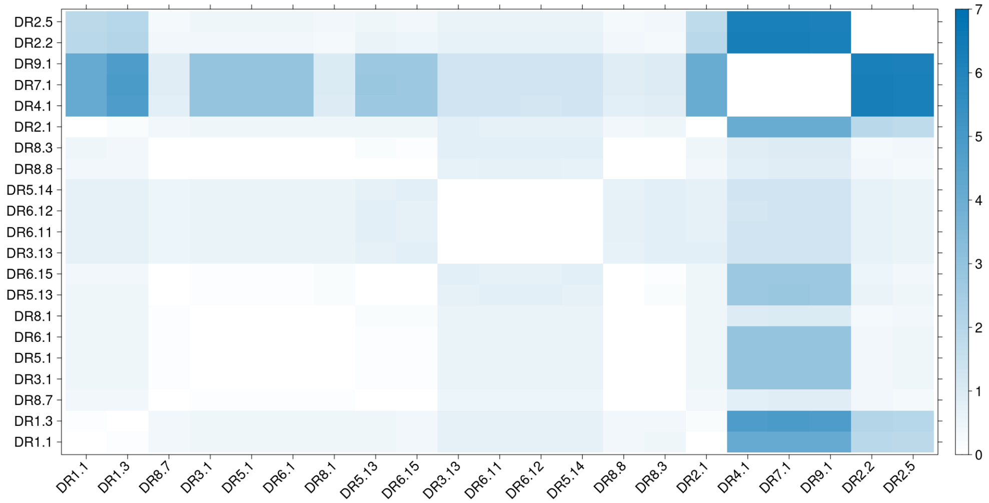


**Figure S7**: **Supplementary Figure 7.** Distance matrix of all HLA-DR-DQ haplotypes. The color bar corresponds to the number of different bases per 100bp.

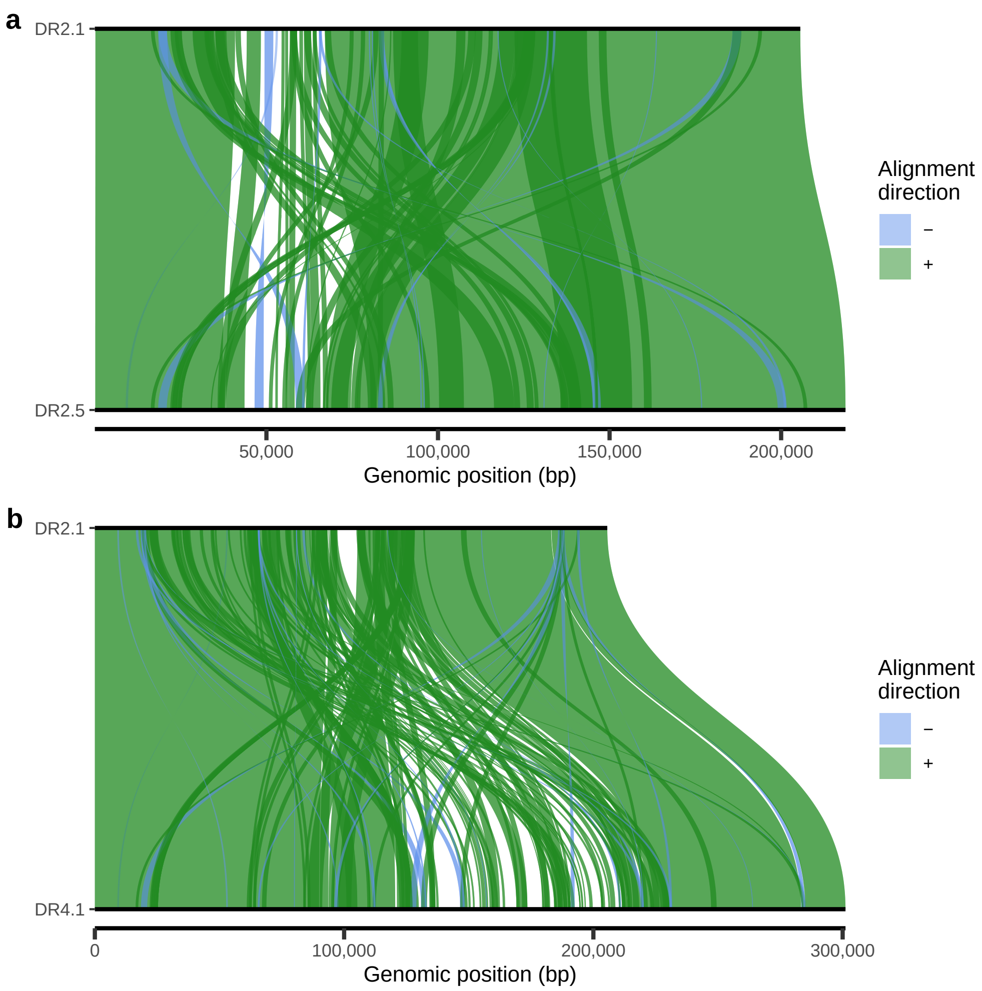


**Figure S8**: **Supplementary Figure 8.** Alignment of DR2.1 with DR2.5 (**a**) and DR4.1 (**b**) shows the possible recombination which created the DR2.1 haplotype. Visualization done with SVbyEye (*108*).

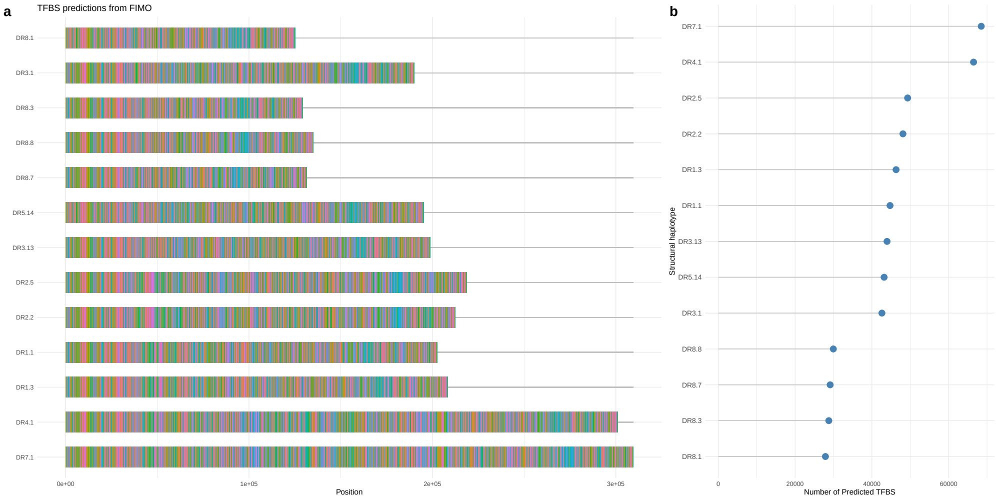


**Figure S9**: **Supplementary Figure 9. a** Transcription factor binding site (TFBS) locations predicted for each structural haplotype using FIMU. Each different color represents a different TFBS motif. **b** Number of predicted TFBSs per structural haplotype.

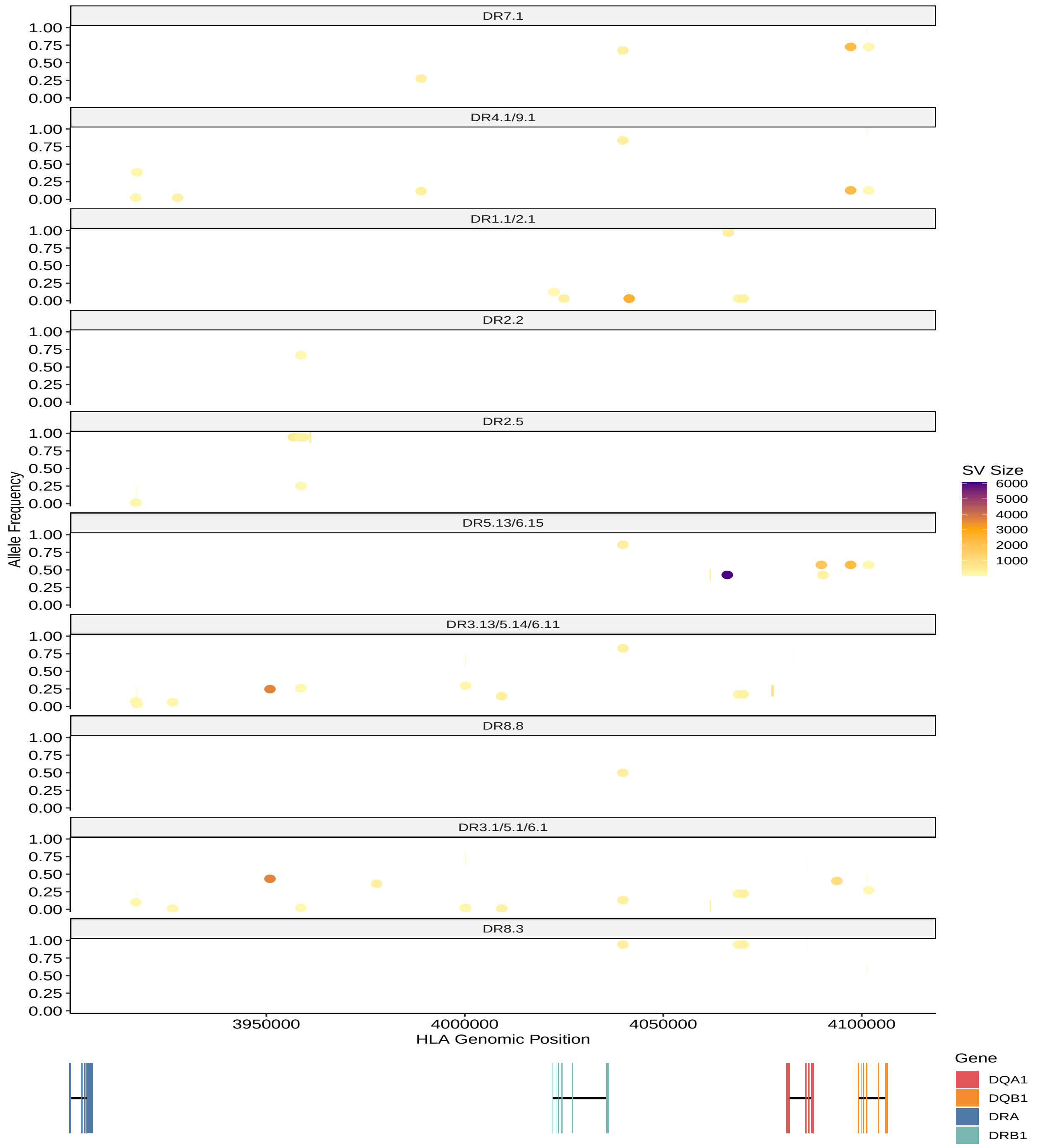


**Figure S10**: **Supplementary Figure 10. Within-haplotype structural diversity across HLA-DR-DQ.** Structural variability within each haplotype across the HLA-DR-DQ region and their respective allelic frequency within each haplotype. The colours represent the size of the SV while dots represent insertions and rectangles deletions, their width informing of their length. The 4 genes with their exon and intronic sequences are also represented, to analyse possible structural variations affecting gene sequences. Genome coordinates relative to the start of the HLA region (Fig.1c).

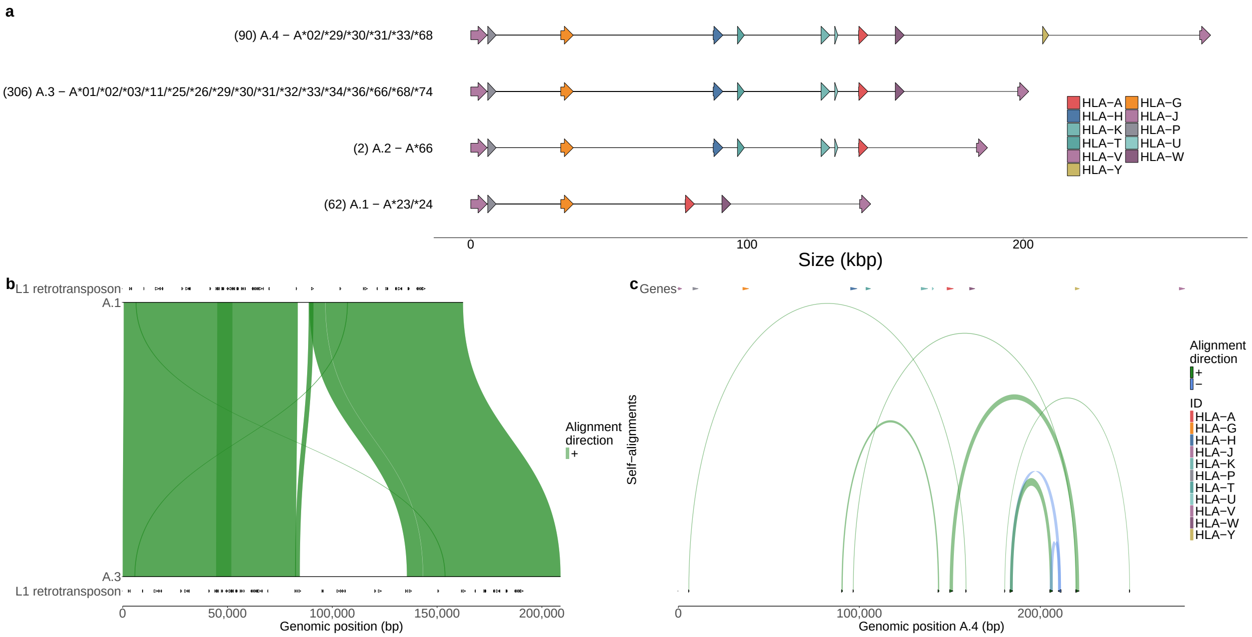


**Figure S11**: **Supplementary Figure 11. Structural diversity across *HLA-A*. a** Four distinct *HLA-A* structural haplotypes identified in 460 haplotypes. The numbers in parentheses indicate the number of haplotypes identified with a specific structure. Each haplotype name is also annotated with all the *HLA-A* alleles identified per-cluster. Haplotypes are ordered by their size which also determines the nomenclature (smaller to larger; 1-4). **b** A pairwise alignment of an A.1 haplotype against an A.3 haplotype with LINE1 retrotransposon annotations. Visualisation done with SVbyEye (*108*). **c** Self-alignment of an A.4 haplotype to visualise possible homologous regions which could explain the deletions observed across structural haplotypes. Visualisation done with SVbyEye.

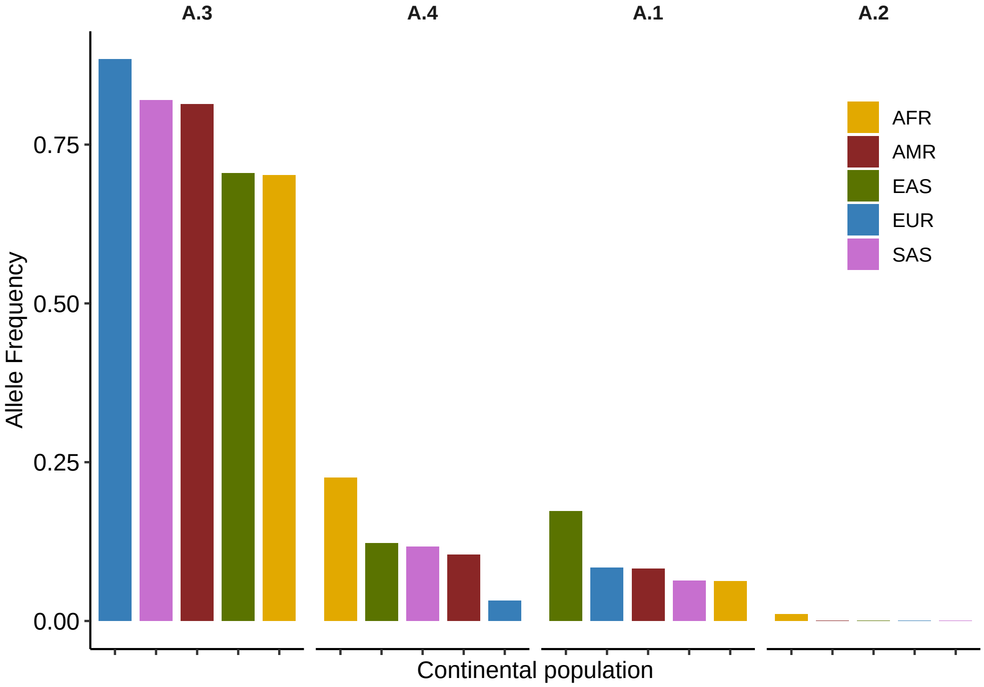


**Figure S12**: **Supplementary Figure 12.** *HLA-A* structural haplotype frequencies across global continental populations.

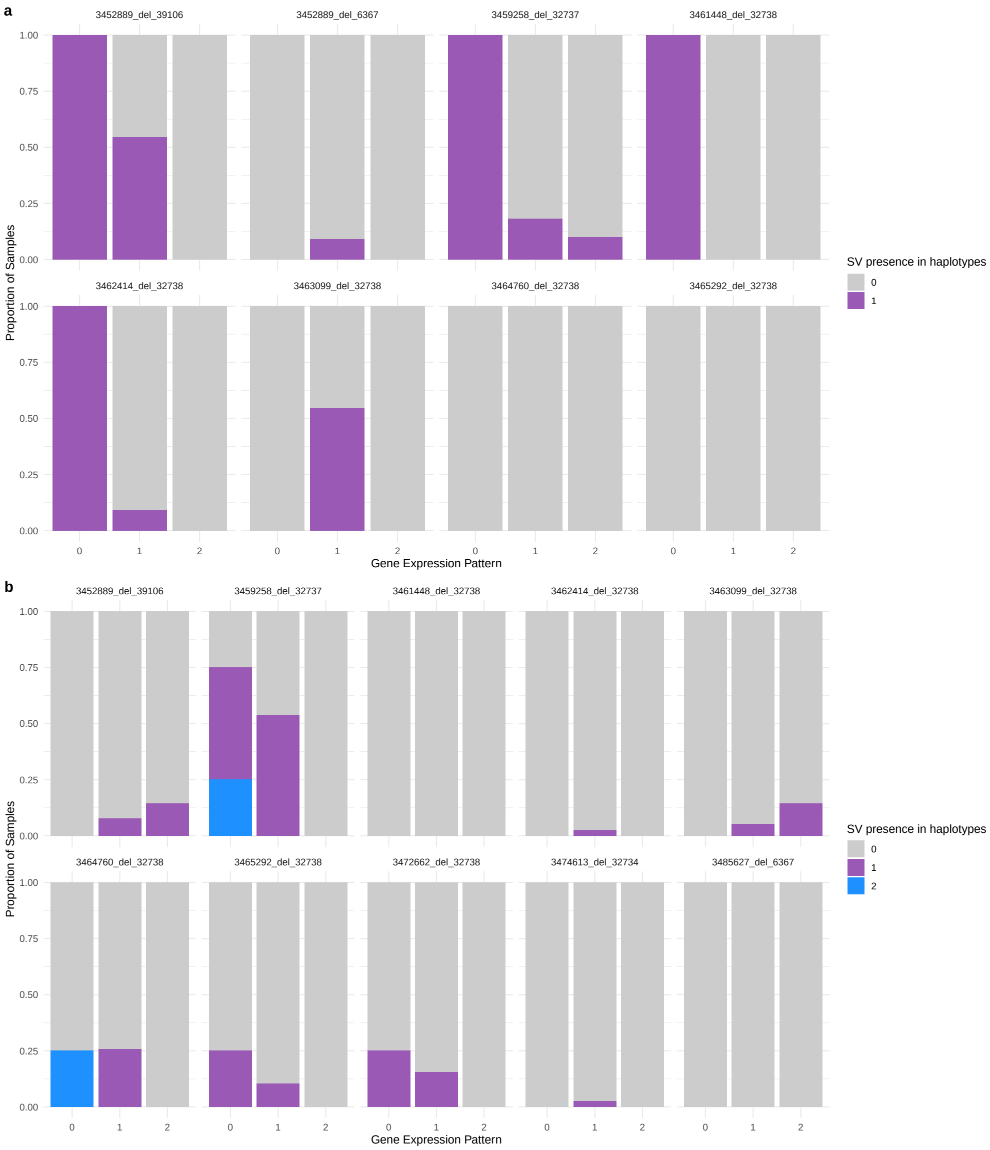


**Figure S13**: **Supplementary Figure 13.** Barplots showing the expression patterns (0 = no expression, 1 = expression in 1 haplotype, 2 = expression in 2 haplotypes) for *C4A* (**a**) and *C4B* (**b**) for each deletion overlapping these genes (0 = no deletion, 1 = deletion in 1 haplotype, 2 = deletion in 2 haplotypes). Samples were filtered out if they had ≥5% unphased reads.

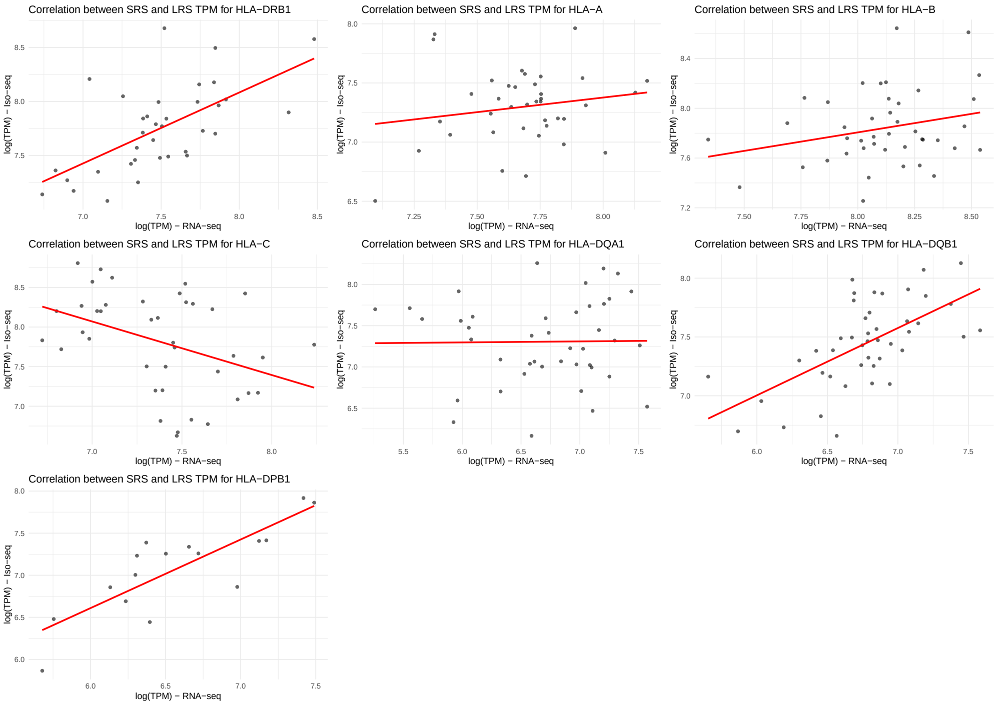


**Figure S14**: **Supplementary Figure 14.** Correlation of HLA gene expression in Iso-seq and RNA-seq datasets in 77 samples across 7 HLA genes. Only samples with ≤5% unphased reads were kept, per gene (see Methods).

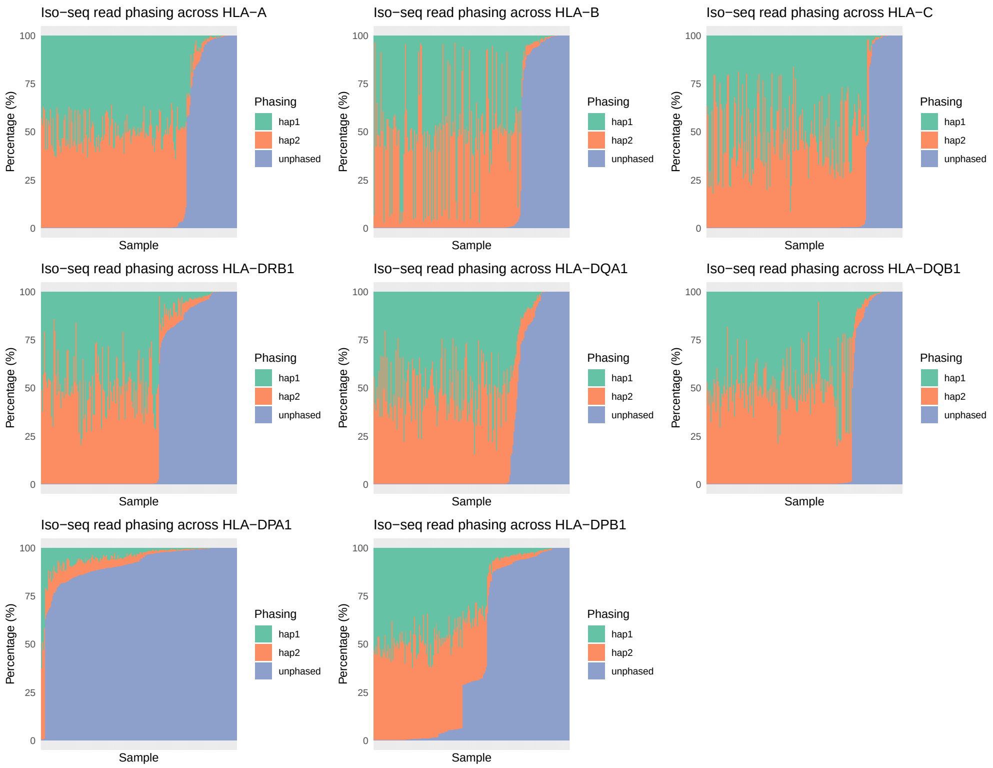


**Figure S15**: **Supplementary Figure 15.** Phasing of Iso-seq reads mapping them onto their respective phased assemblies across their HLA genes.

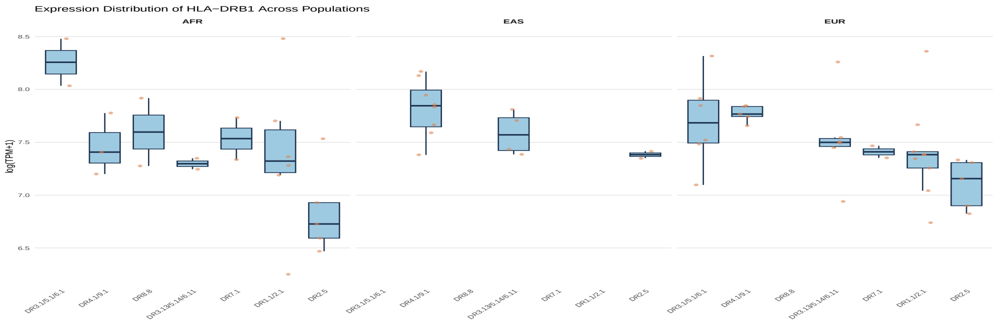


**Figure S16**: **Supplementary Figure 16.** *HLA-DRB1* expression in each DR-DQ structural haplotype across all continental populations as boxplots. Box plots indicate the median (center line), interquartile range (box), and whiskers extending to 1.5 × the interquartile range.

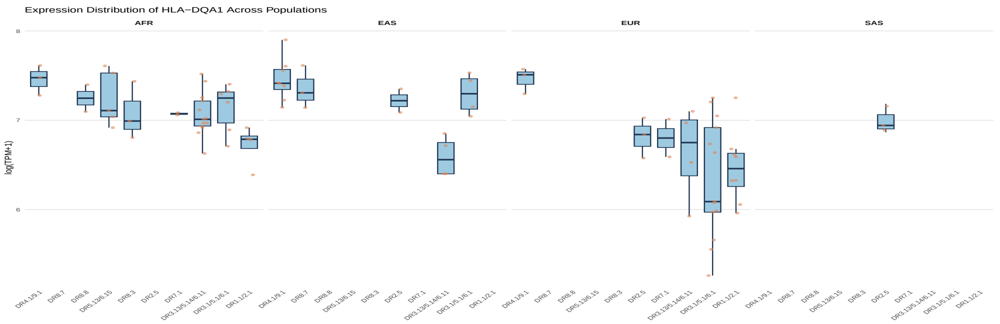


**Figure S17**: **Supplementary Figure 17.** *HLA-DQA1* expression in each DR-DQ structural haplotype across all continental populations as boxplots. Box plots indicate the median (center line), interquartile range (box), and whiskers extending to 1.5 × the interquartile range.

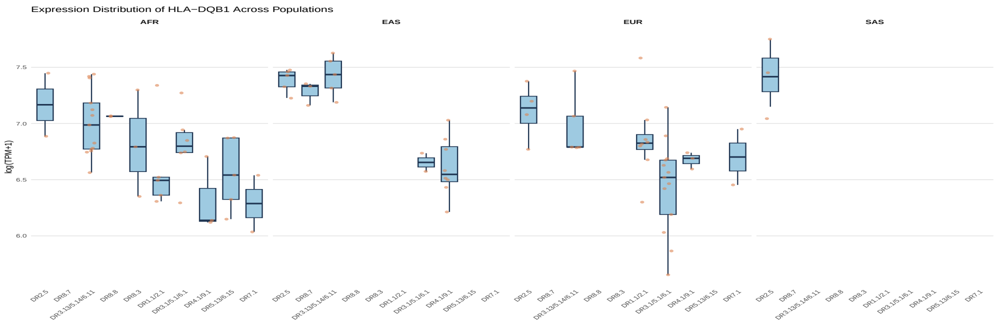


**Figure S18**: **Supplementary Figure 18.** *HLA-DQB1* expression in each DR-DQ structural haplotype across all continental populations as boxplots. Box plots indicate the median (center line), interquartile range (box), and whiskers extending to 1.5 × the interquartile range.

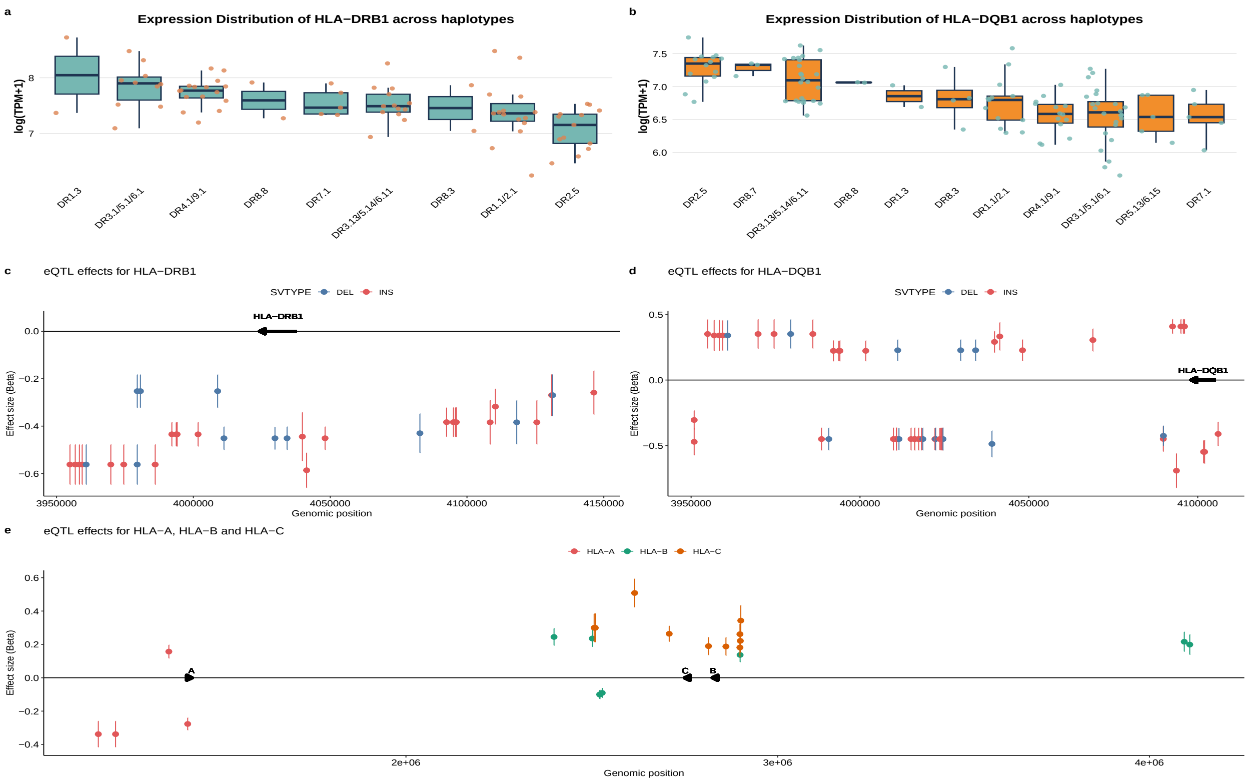


**Figure S19**: **Supplementary Figure 19. HLA SV-eQTLs**. **a** *HLA-DRB1* expression across HLA-DR-DQ haplotypes. Box plots indicate the median (center line), interquartile range (box), and whiskers extending to 1.5 × the interquartile range. **b** *HLA-DQB1* expression across HLA-DR-DQ haplotypes. Box plots indicate the median (center line), interquartile range (box), and whiskers extending to 1.5 × the interquartile range. **c** Position and effect size of significant SV-eQTLs associated with modified *HLA-DRB1* gene expression with standard error bars. **d** Position and effect size of significant SV-eQTLs associated with modified *HLA-DQB1* gene expression with standard error bars. **e** Position and effect size of SV-eQTLs associated with modified *HLA-A*, *HLA-B* and *HLA-C* gene expression with standard error bars.

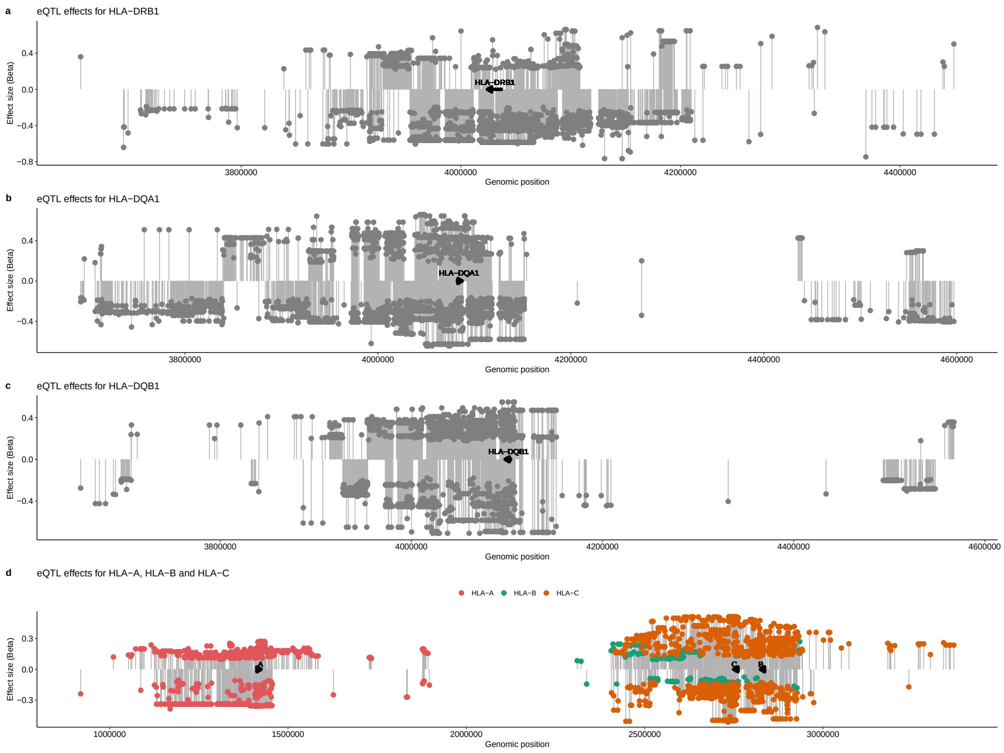


**Figure S20**: **Supplementary Figure 20.** Position and effect size of SNP-eQTLs associated with modified *HLA-DRB1* (**a**), *HLA-DQA1* (**b**), *HLA-DQB1* (**c**) and *HLA-A*, *-B* and *-C* (**d**) gene expression.

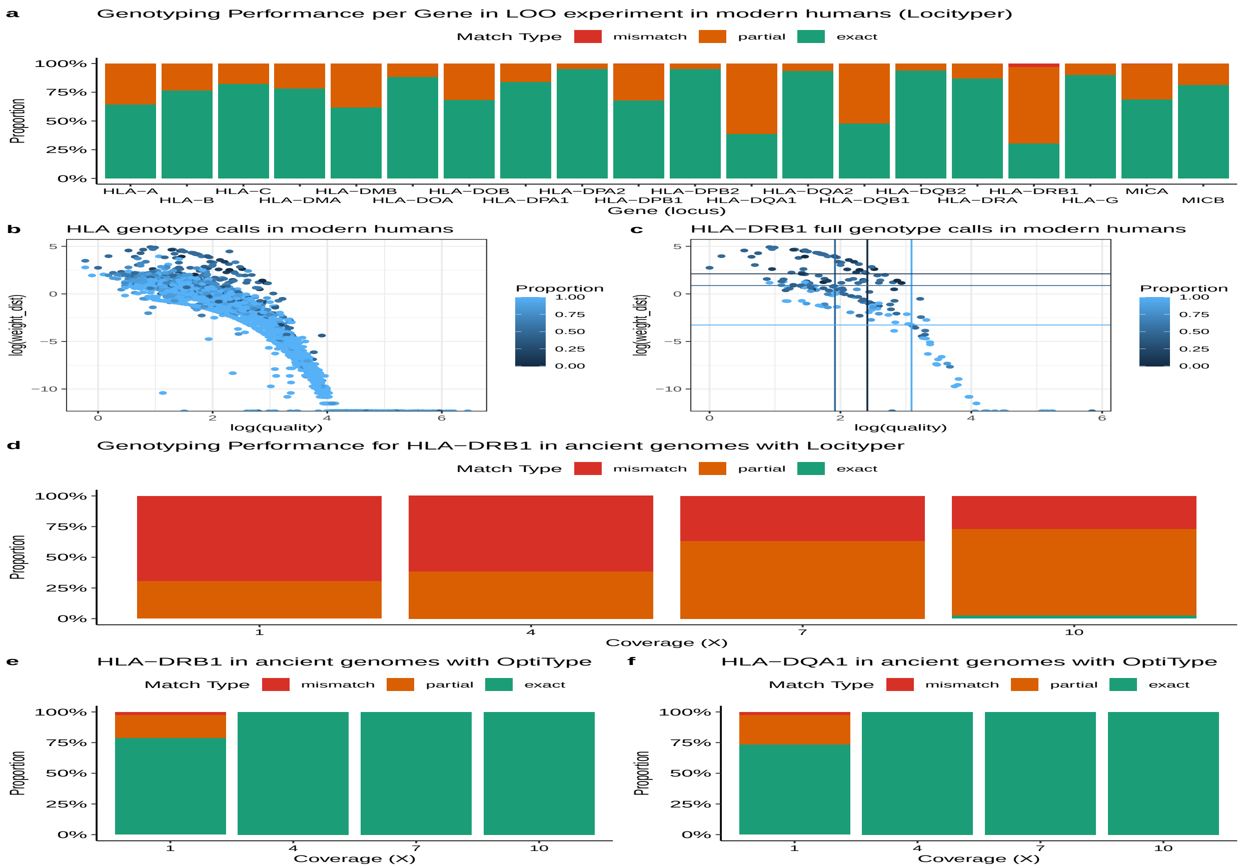


**Figure S21**: **Extended Data Figure 21. Modern and ancient human HLA genotyping benchmark.** **a** Benchmark of HLA gene typing on modern human short-read data with a leave-one-out approach using Locityper (458 haplotypes in the Locityper database). Partial matches can range from a single difference in the 4th field to differences across most fields. **b** Log transformation of weighed distance and quality of all genotype calls, across all genes. The color of each genotype corresponds to the proportion of correct calls. A proportion score of 1 means an exact match in both genotypes, scores between 0-1 mean different matches across fields, with higher scores implying smaller differences in the 4th or 3rd fields, while 0 implies a complete mismatch. **c** Same plot as **b** for *HLA-DRB1*. Horizontal and vertical lines show the median values for weighed distances and quality scores, respectively, across exact, partial, and mismatches. **d** Benchmark of the first field in *HLA-DRB1* typing across various coverages in simulated ancient samples using Locityper. **e** Benchmark of the first field in *HLA-DRB1* typing across various coverages in simulated ancient samples using OptiType. **f** Benchmark of the first field in *HLA-DQA1* typing across various coverages in simulated ancient samples using OptiType.

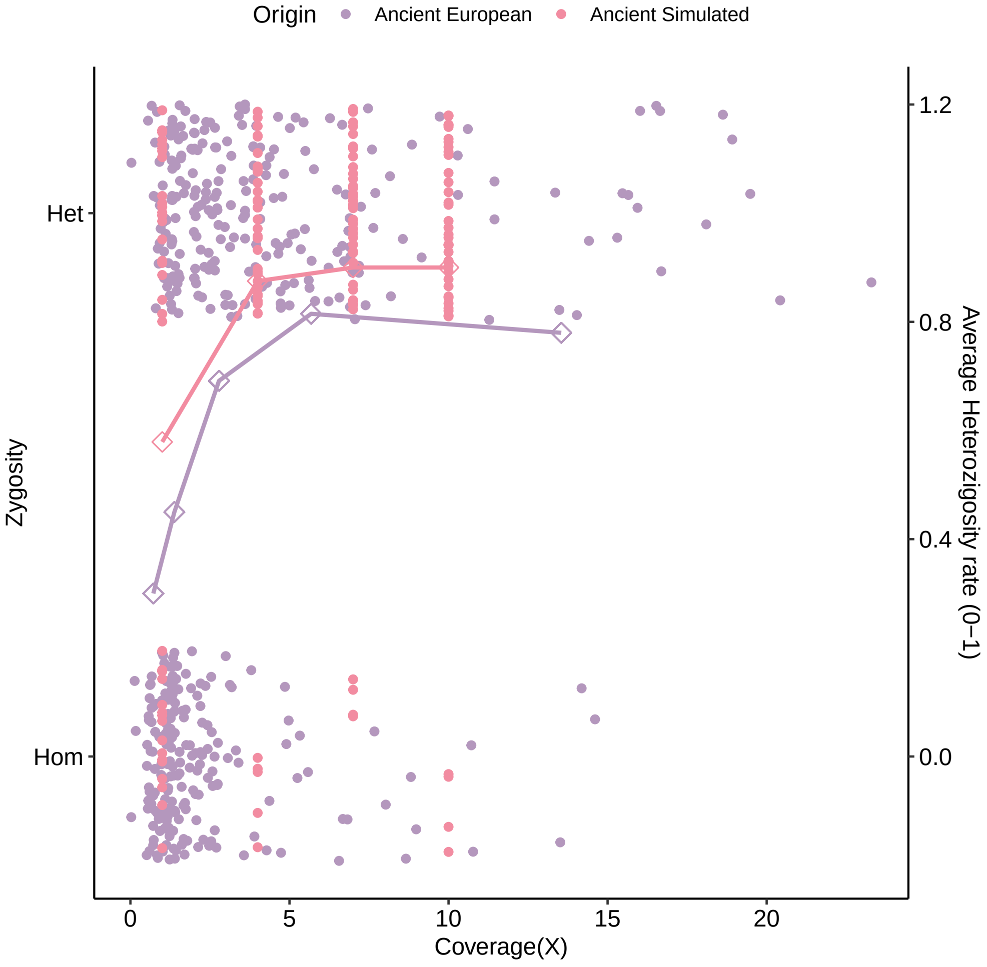


**Figure S22**: **Supplementary Figure 22.** Heterozigosity rates across diverse coverages of European and simulated ancient DNA samples. Rhomboid shapes represent the average heterozigosity frequency across bins [0, 1], (1, 2],(2, 4], (4, 8], (8, 25] for ancient European and 1, 4, 7 and 10 for ancient simulated.

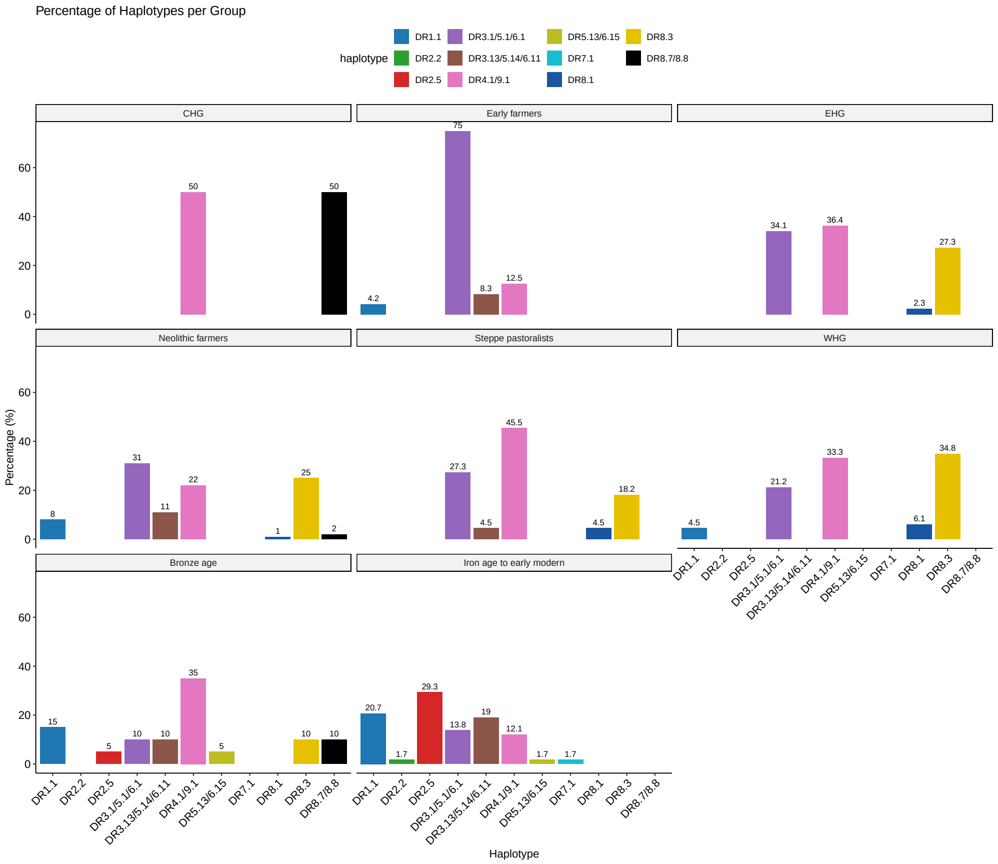


**Figure S23**: **Supplementary Figure 23.** Structural haplotype frequencies across ancient European populations. CHG: Caucasus hunter-gatherer; EHG: Eastern hunter-gatherer; WHG: Western hunter-gatherer.

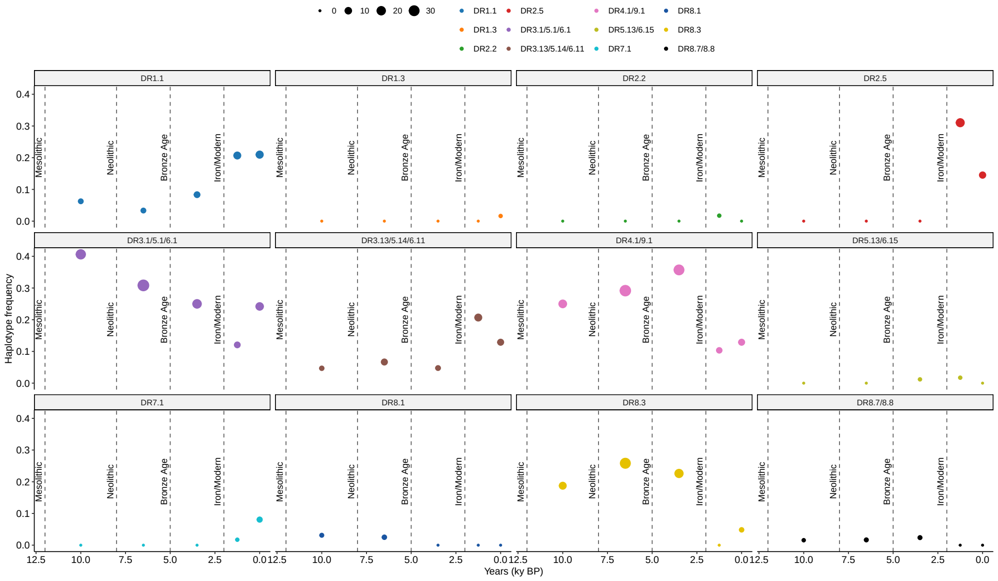


**Figure S24**: **Supplementary Figure 24.** Frequency of structural haplotypes over time. Time bins (unit in kyr BP): [12,8), [8, 5), [5, 2), [2, 0.5), [0.5, 0]. The points represent the average within each bin, with shaded lines representing standard errors.

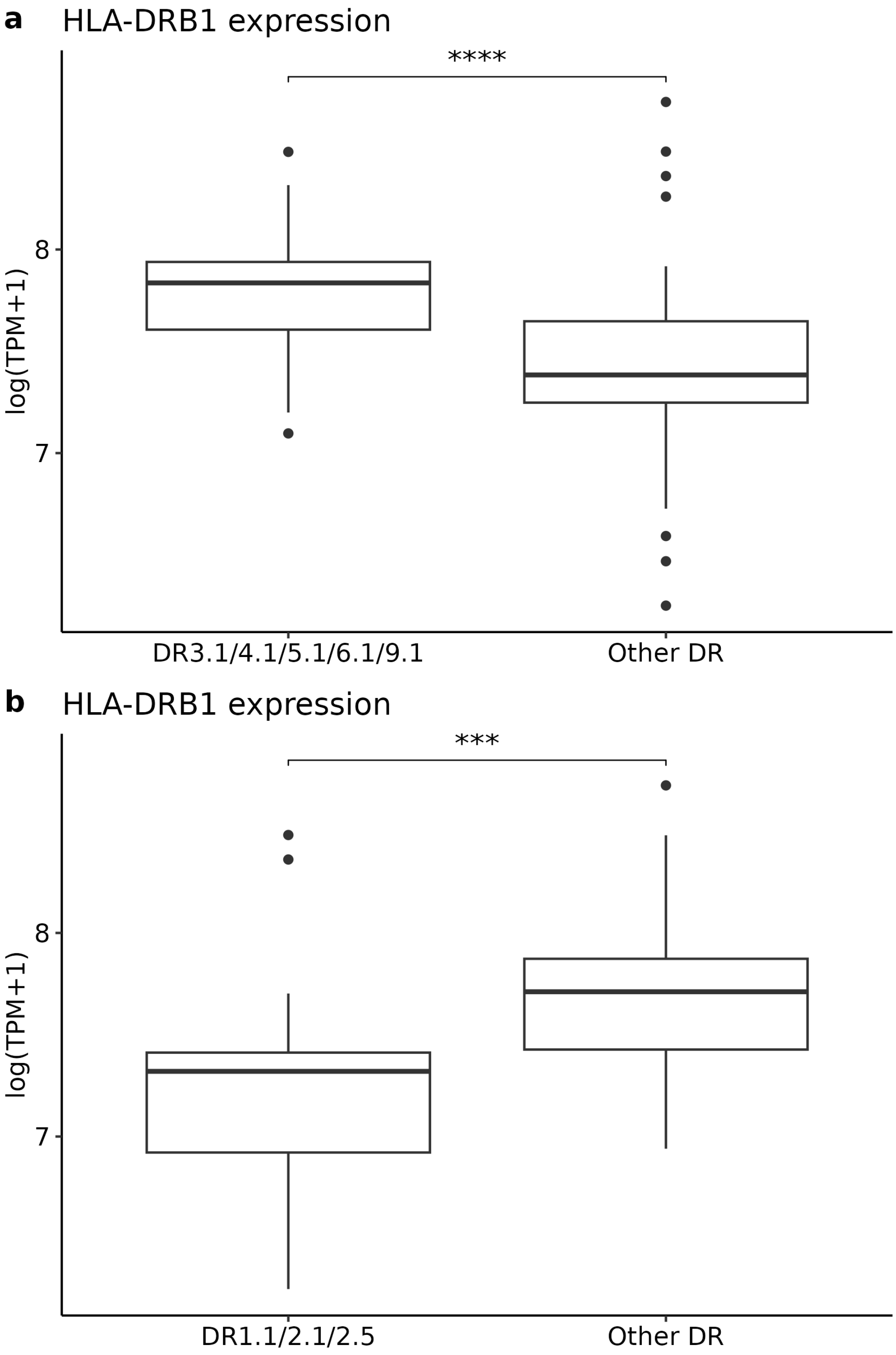


**Figure S25**: **Supplementary Figure 25.** Significant differences in high (**a**) and low (**b**) *DRB1*-expressing structural haplotypes. Box plots indicate the median (center line), interquartile range (box), and whiskers extending to 1.5 × the interquartile range.

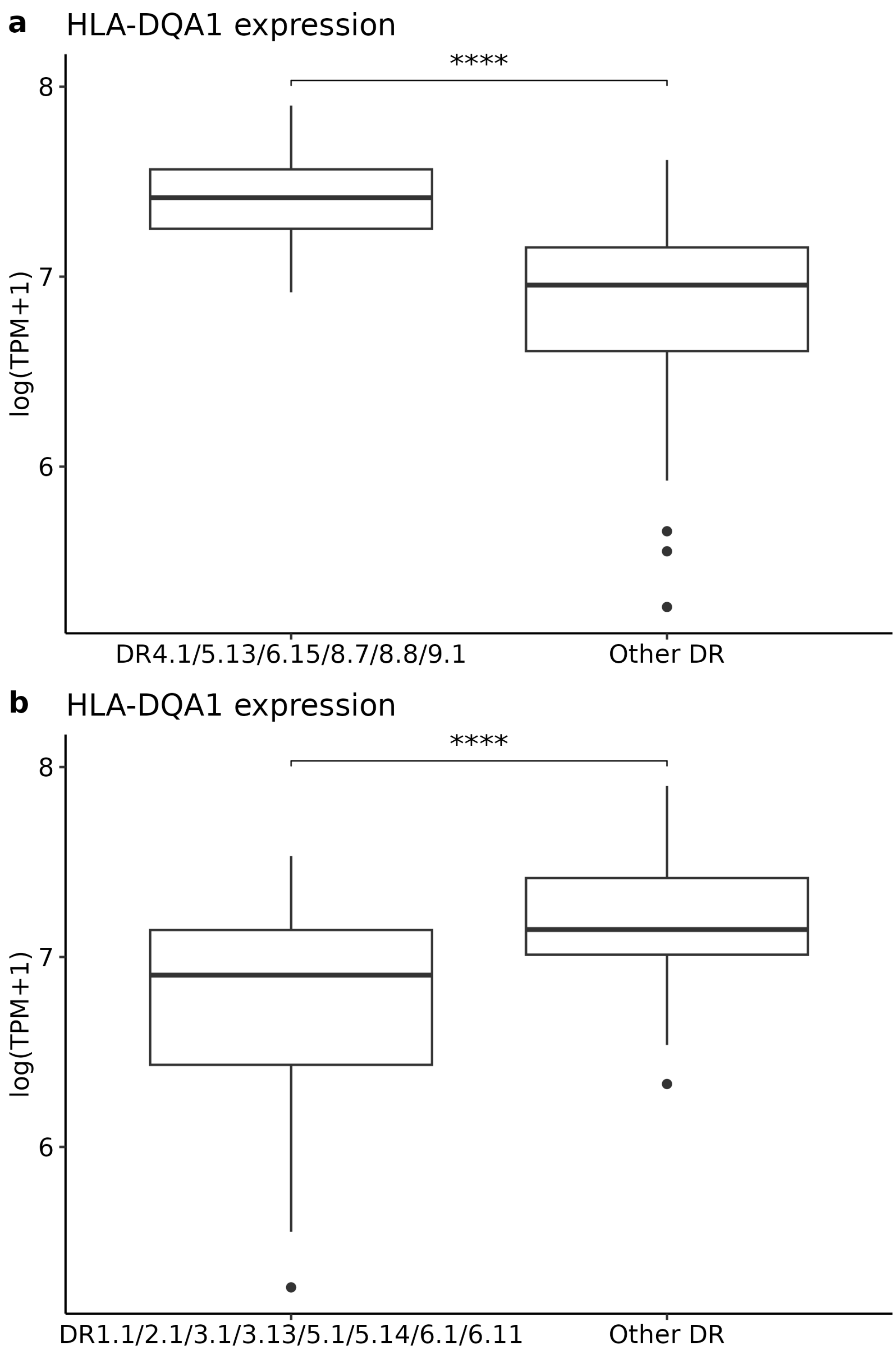


**Figure S26**: **Supplementary Figure 26.** Significant differences in high (**a**) and low (**b**) *DQA1*-expressing structural haplotypes. Box plots indicate the median (center line), interquartile range (box), and whiskers extending to 1.5 × the interquartile range.

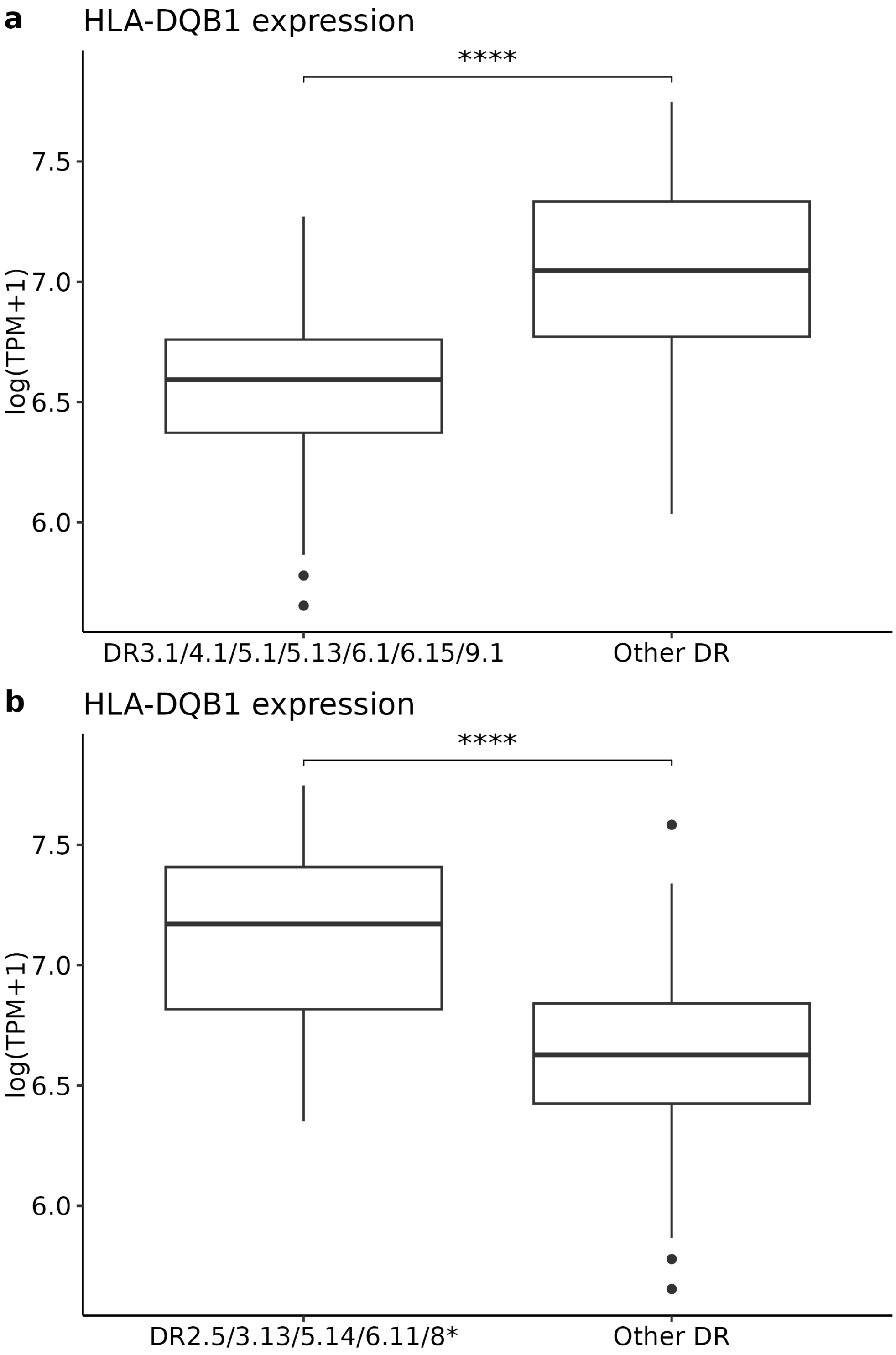


**Figure S27**: **Supplementary Figure 27.** Significant differences in low (**a**) and high (**b**) *DQB1*-expressing structural haplotypes. Box plots indicate the median (center line), interquartile range (box), and whiskers extending to 1.5 × the interquartile range.

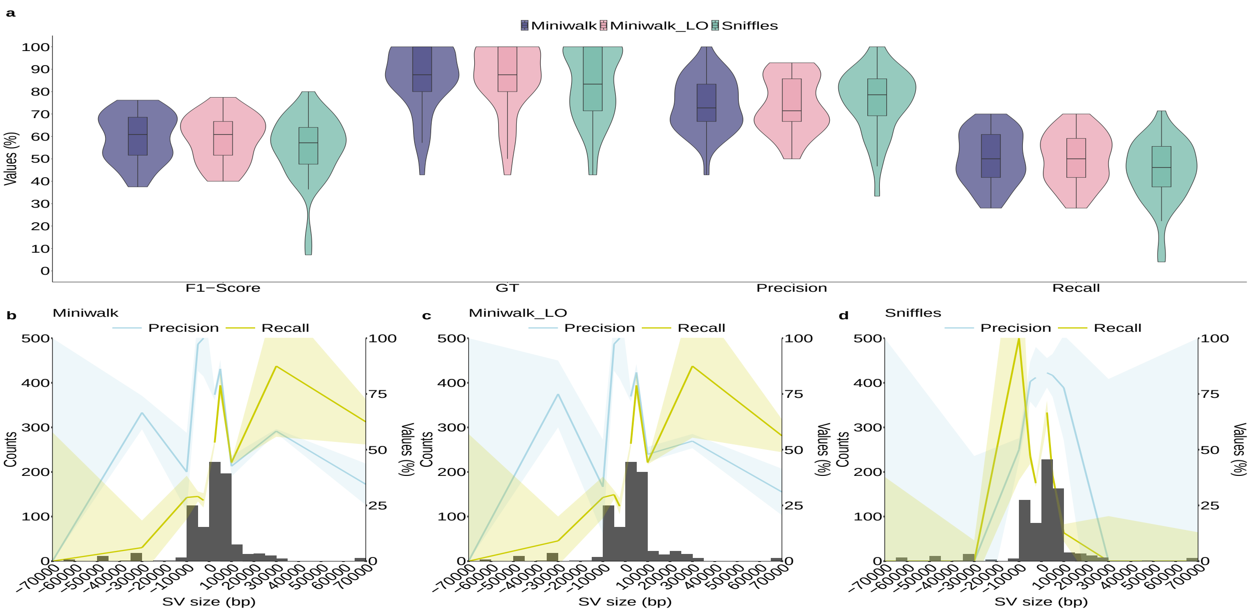


**Figure S28**: **Supplementary Figure 28. Long-read genotyping of large (≥2,500 bp) SVs across the HLA benchmark. a** Benchmark of large (≥2,500 bp) SV genotyping with long reads (ONT (*20*))using miniwalk with the full HLA pangenome graph, leaving out the testing samples from the graph (Minigraph_LO) and Sniffles. Each data point represents an SV call set from one assembly, with assemblies serving as independent biological replicates. No technical replicates were used. No explicit control group was included; instead, distributions reflect genotyping performance across the independent assemblies. Box plots indicate the median (centre line), interquartile range (box), and whiskers extending to 1.5× the interquartile range. GT: genotype concordance. **b-d** Histogram with 5,000 bp bins showing the found SVs in **a**, with two segments with 2,500 bp bin from SVs of size 2,500-5,000 bp, 5,000 bp bin for SVs of size 5,000-10,000 bp, 20,000 bp bin for SVs of size 10,000-30,000 bp, 40,000 bp bin for SVs of size 30,000-70,000 bp and SVs larger than 70,000 bp grouped together. The two segments show the precision and recall means across different SV sizes. Shaded regions represent 95% confidence intervals (mean ± 1.96 × standard error).

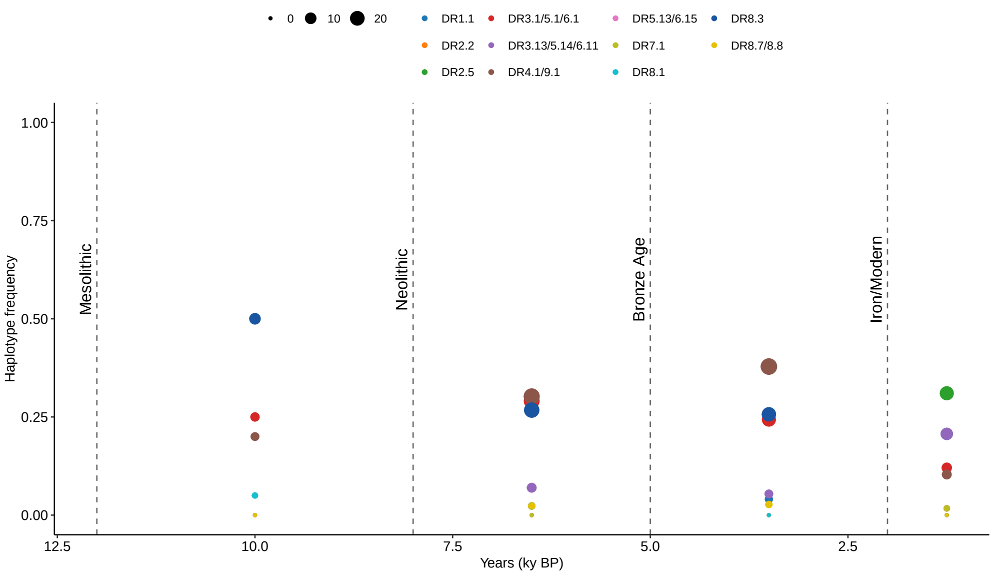


**Figure S29**: **Supplementary Figure 29.** Frequency of structural haplotypes over time, in Northern Europe. These trends show similar patterns when including the whole dataset, proving that the haplotype frequencies were similar across northern and southern Europe. Time bins (unit in kyr BP): [12,8), [8, 5), [5, 2), [2, 0.5), [0.5, 0]. The points represent the average within each bin.

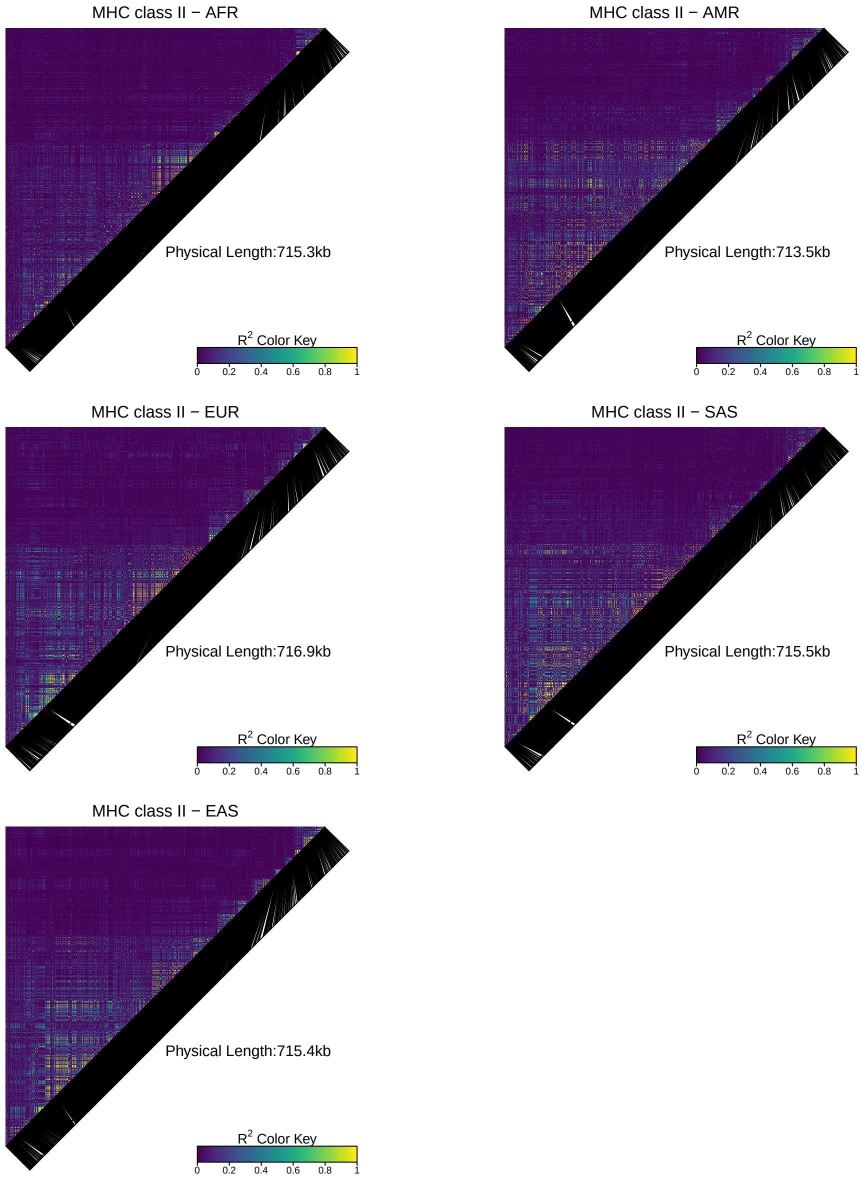


**Figure S30**: **Supplementary Figure 30.** Linkage disequilibrium in each human population across 700 kbp of MHC class II from *HLA-DRA* to *HLA-DPB2*. The large LD block spans DRA-DQB1 across all populations.

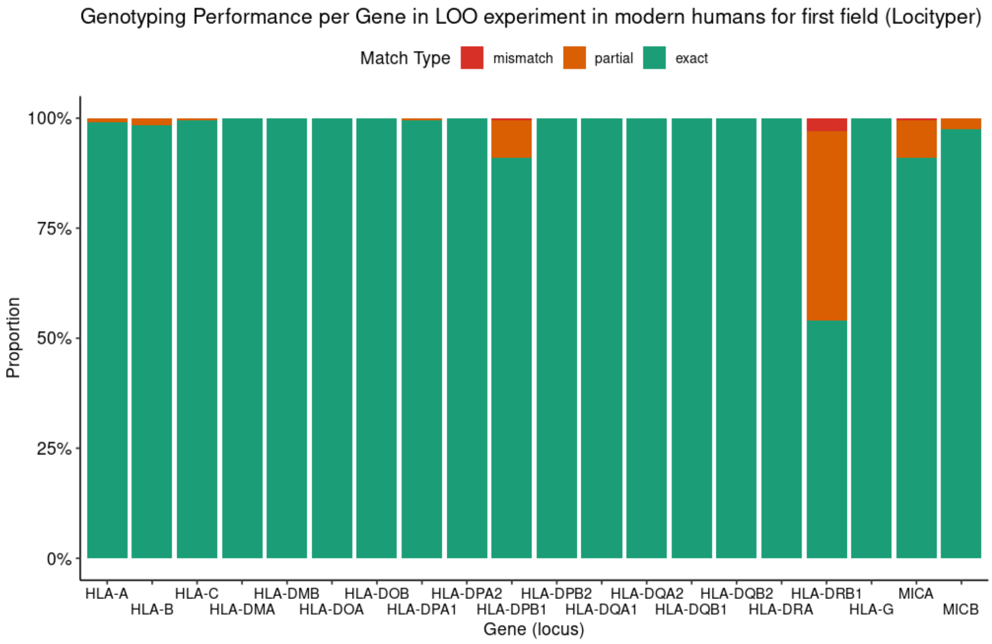


**Figure S31**: **Supplementary Figure 31.** Benchmark of HLA gene typing on modern human short-read data with a leave-one-out approach. For this benchmark, only the first field in the typing scheme was bench marked. Partial matches refer to those where only one of two genotypes are correctly predicted.

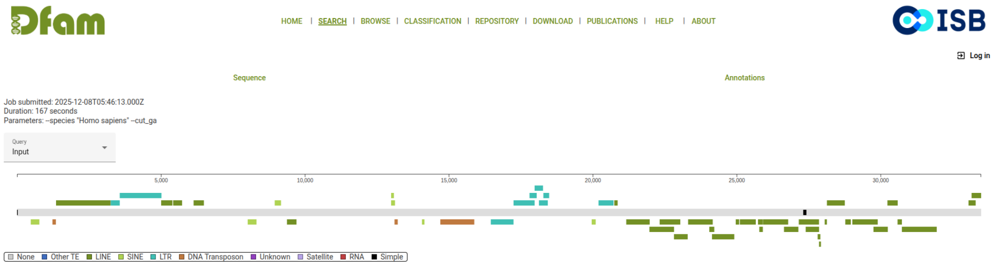


**Figure S32**: **Supplementary Figure 32.** Dfam repetitive sequences mapped to the first 30kbps of sequence from a 71kbp false positive SV lowering the precision in the SV benchmark.
